# Accurate and Complete Genomes from Metagenomes

**DOI:** 10.1101/808410

**Authors:** Lin-Xing Chen, Karthik Anantharaman, Alon Shaiber, A. Murat Eren, Jillian F. Banfield

## Abstract

Genomes are an integral component of the biological information about an organism and, logically, the more complete the genome, the more informative it is. Historically, bacterial and archaeal genomes were reconstructed from pure (monoclonal) cultures and the first reported sequences were manually curated to completion. However, the bottleneck imposed by the requirement for isolates precluded genomic insights for the vast majority of microbial life. Shotgun sequencing of microbial communities, referred to initially as community genomics and subsequently as genome-resolved metagenomics, can circumvent this limitation by obtaining metagenome-assembled genomes (MAGs), but gaps, local assembly errors, chimeras and contamination by fragments from other genomes limit the value of these genomes. Here, we discuss genome curation to improve and in some cases achieve complete (circularized, no gaps) MAGs (CMAGs). To date, few CMAGs have been generated, although notably some are from very complex systems such as soil and sediment. Through analysis of ~7000 published complete bacterial isolate genomes, we verify the value of cumulative GC skew in combination with other metrics to establish bacterial genome sequence accuracy. Interestingly, analysis of cumulative GC skew identified potential mis-assemblies in some reference genomes of isolated bacteria and the repeat sequences that likely gave rise to them. We discuss methods that could be implemented in bioinformatic approaches for curation to ensure that metabolic and evolutionary analyses can be based on very high-quality genomes.

## Introduction

In an opinion paper published relatively early in the microbial genomics era, Fraser et al. stated “you get what you pay for” (Fraser et al. 2002). The authors argued the lower scientific value of draft (partial) vs. complete genomes, noting for example higher error rates, potential contaminant sequences, loss of information about gene order, lower ability to distinguish additional chromosomes and plasmids, and most importantly, missing genes. Despite the clarity of this view, the field moved toward the generation of draft isolate genomes to optimize the rate of supply of new sequence information and to lower the cost. Genome-resolved metagenomics has almost exclusively settled for uncurated draft genomes, now often referred to as metagenome-assembled genomes (MAGs). A summary of the basic methods for generating MAGs was provided by (Sangwan et al. 2016). A more recent review provides an overview of assembly methods and offers some insights into the complexity of genome recovery from metagenomes and a valuable overview of certain types of assembly errors that can occur (Olson et al. 2017).

The first MAGs were published in 2004 (Tyson et al. 2004) and there are now hundreds of thousands of them in public databases. The ever increasing depth of high-throughput sequencing now make even the most challenging environments with low archaeal, bacterial, and viral biomass, such as insect ovaries (Reveillaud et al. 2019), human gut tissue biopsies (Vineis et al. 2016), hospital room surfaces (Brooks et al. 2017), and even human blood (Moustafa et al. 2017) amenable to shotgun metagenomic surveys and recovery of MAGs. Although incomplete, draft MAGs represent a major advance over knowing nothing about the genes and pathways present in an organism, and led to the discovery of new metabolisms. For example, the complete oxidation of ammonia to nitrate via nitrite (i.e., comammox) was determined by the detection of necessary genes in a single MAG (Daims et al. 2015; van Kessel et al. 2015). MAGs are often derived from uncultivated organisms that can be quite distantly related to any isolated species, which is a clear advantage of MAGs (Becraft et al. 2017; Garg et al. 2019). For this reason, genome-resolved metagenomics has been critical for more comprehensive descriptions of bacterial and archaeal diversity and the overall topology of the Tree of Life (Hug et al. 2016).

Counter to this view, there is some sentiment that MAGs are not useful because they are composites and thus not representative of their populations (Becraft et al. 2017). However, a genome reconstructed from a clonal microbial culture also does not represent the cloud of biologically important variation that exists in the natural population from where the isolate was derived. Population diversity can be analyzed by comparing all individual sequences (or short reads) to the metagenome-assembled reference genome (Simmons et al. 2008; Delmont et al. 2019). While some populations are near-clonal, others are very complex strain mixtures and yet others fall on the continuum between these (Lo et al. 2007; Chivian et al. 2008; Simmons et al. 2008). As strain divergence leads to assembly fragmentation (expanded on below), high quality genomes are unlikely to be generated for relatively heterogeneous populations. Assembly of exceptionally long fragments from short read (e.g., Illumina) data is only anticipated when within population diversity is low, as may occur following a recent bloom, selective sweep, or due to recent colonization by a single cell or a small cluster of closely related cells. In such cases, the genomes that assemble well are typically highly representative of the population from which they are derived and the vast majority of reads report the same base at the same position. For example, in one recently published complete 4.55 Mbp genome (Banfield et al. 2017), the frequency of single nucleotide variants (SNVs) is ~ 0.12% (Figure 1), not substantially different from the expected sequencing error rate (0.04-0.12%) (Schirmer et al. 2016).

**Figure 1.**
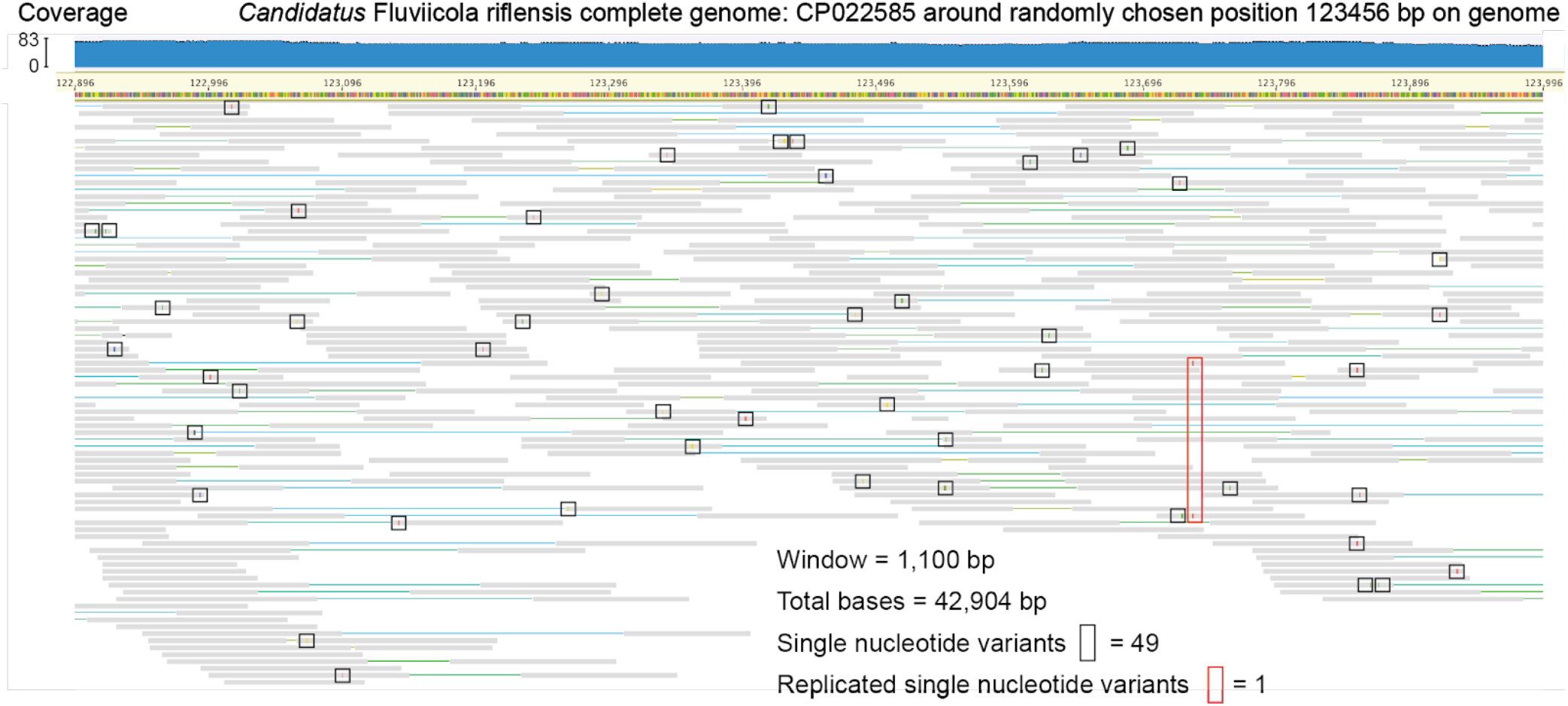
The low frequency of single nucleotide variants (SNVs) of a recently published CMAG. A randomly chosen region, centered on position 123,456 (1100 bp in length) of the CMAG of *Candidatus* Fluviicola riflensis is shown with mapped reads (Banfield et al. 2017). SNVs that only occur once are indicated by black boxes and the one replicated SNV indicated by a red box. Clearly, the consensus sequence is well supported.

### Assembly and binning are important steps in metagenomic studies

Assembly of short metagenomic reads into contiguous segments of DNA is a computationally intensive task, and its effectiveness often depends on the complexity of the environment (Sharon and Banfield 2013). However, assembly of contigs/scaffolds offers many advantages over short-read based analyses. First, they enable the identification of complete open reading frames. Second, assemblies provide larger genomic contexts (e.g., operons). In combination, these considerations improve predictions of metabolic capacities. Further, assembled sequences provide information about gene synteny and better resolve taxonomic profiles (e.g., by providing sets of proteins for taxonomy based on concatenated proteins encoded in the same genome (e.g., (Hug et al. 2016; Parks et al. 2018)). These improvements can overcome misleading interpretations of short-read data (Ackelsberg et al. 2015; Afshinnekoo et al. 2015).

The critical step required to establish a genome from a metagenomic assembly is binning. This involves assignment of assembled fragments to a draft genome based on detection on any scaffold of some signal(s) that occur(s) locally within a genome and persists genome-wide. Most commonly used features that can facilitate accurate binning of scaffolds include depth of sequencing measured by read coverage, sequence composition measured for example by tetranucleotide composition, and phylogenetic profile measured by the ‘best taxonomic hits’ for each predicted protein on each scaffold. Sometimes, and mostly in datasets from very simple communities or for highly abundant organisms, the process of binning can be as easy as collecting together all fragments that share a single clearly defined feature (Figure S1), such as a discrete set of scaffolds with similar coverage, or unique and well-defined tetranucleotide patterns or GC content. In other cases, a combination of a few well defined signals, such as GC content, coverage, and phylogenetic profile of scaffolds, are sufficient to clearly define a bin. However, over reliance on phylogenetic profile can be dangerous, especially if the genome is for an organism that is only distantly related to those in the databases used for profiling. Further, some fragments can have an unexpected phylogenetic profile relative to the rest of the genome because the region has not been encountered previously in genomes of related organisms, possibly because it was acquired by lateral gene transfer. Thus, the most robust bins will draw on a combination of multiple clear signals.

If a study includes a set of samples with related community membership, an important constraint for bin assignment can be provided by the shared patterns of abundance of a fragment across a sample series. The use of series samples data for binning was first proposed by Sharon et al. (Sharon et al. 2013), and this strategy is now a central feature in most automated binning algorithms, including CONCOCT (Alneberg et al. 2013), MaxBin (Wu et al. 2014), ABAWACA (Brown et al. 2015) and MetaBAT (Kang et al. 2015), as well as manual binning and MAG refinement strategies (Wrighton et al. 2012; Shaiber and Eren 2019). Series based binning can exclude contaminant scaffolds from a MAG whose abundance shows a different pattern over time/space/treatment. We have found that no single binning algorithm is the most effective for all sample/environment types or even for all populations within one sample. The recently published method DASTool tests a flexible number of different binning methods, evaluates all outcomes and chooses the best bin for each population (Sieber et al. 2018). A similar strategy has been utilized in a modular pipeline software called MetaWRAP (Uritskiy et al. 2018).

### A case study: Binning can greatly improve data interpretation

Contigs that do not represent entire chromosomes may not be appropriate proxies for microbial populations without binning, and claims made based on unbinned contigs can lead to erroneous conclusions. For instance, a recent study focusing on human blood used shotgun metagenomic sequencing of circulating cell-free DNA from more than 1,000 samples and recovered a large number of contigs with novel bacterial and viral signatures (Kowarsky et al. 2017), suggesting that “hundreds of new bacteria and viruses” were present in human blood, and that this environment contained more microbial diversity than previously thought. While the authors performed PCR experiments to independently confirm the existence of some of these signatures in blood samples, they did not attempt to assign assembled contigs to genome bins. Here, we studied contigs from these blood metagenomes with a genome-resolved strategy to investigate the presence of previously unknown bacterial populations.

To explore the origin of bacterial signatures found in the novel set of contigs recovered from cell-free DNA blood metagenomes (Figure 2a), we first searched for the 139 bacterial single-copy core genes (SCGs) described by Campbell et al. (Campbell et al. 2013). This analysis identified 76 bacterial SCGs among all contigs, and of these, 56 occurred only once, suggesting that a single microbial population may explain a large fraction of the bacterial signal found among novel contigs (Figure 2b). Of the 56 genes that occurred only once, 18 were ribosomal proteins. Comparison of the amino acid sequences of these ribosomal proteins to those in the NCBI’s non-redundant protein sequence database revealed that the vast majority of them best matched to proteins from genomes that fall within the recently described ‘Candidate Phyla Radiation’ (CPR) (Brown et al. 2015), a group of microbes with rather small genomes, reduced metabolic capacities (Rinke et al. 2013; Brown et al. 2015), and at least in some cases very small cell sizes (Luef et al. 2015), which suggest largely symbiotic lifestyles (He et al. 2015; Nelson and Stegen 2015). Even though ribosomal proteins found in blood metagenomes best matched to CPR genomes, the levels of sequence identity of these matches were very low, and taxonomic affiliations of best hits were divergent within the CPR (Table S1), which could simply reflect the novelty of a single population rather than the presence of multiple populations. To investigate the distribution of these proteins we clustered novel contigs based on their tetranucleotide frequencies (Figure 2a). We found that most bacterial SCGs occurred in a relatively small group of contigs with similar tetranucleotide composition. Manual selection of these contigs, and their further refinement using additional ‘non-novel’ contigs that were not included in the original study by Kowarsky et al. (2017) resulted in a single CPR MAG that is 613.5 kbp in size with a completion estimate of 52.5%. Our phylogenomic analysis affiliated this MAG with the superphylum Parcubacteria (previously OD1) of the CPR (Figure 2c). Regardless of the origins of this population in these metagenomes, our genome-resolved analysis contrasts with the prior interpretation of these data and suggests that Parcubacteria appears to be the only major novel bacterial group whose DNA is present in human blood metagenomes. This finding demonstrates the critical importance of binning-based strategies to justify claims of microbial diversity in metagenomic analyses.

**Figure 2.**
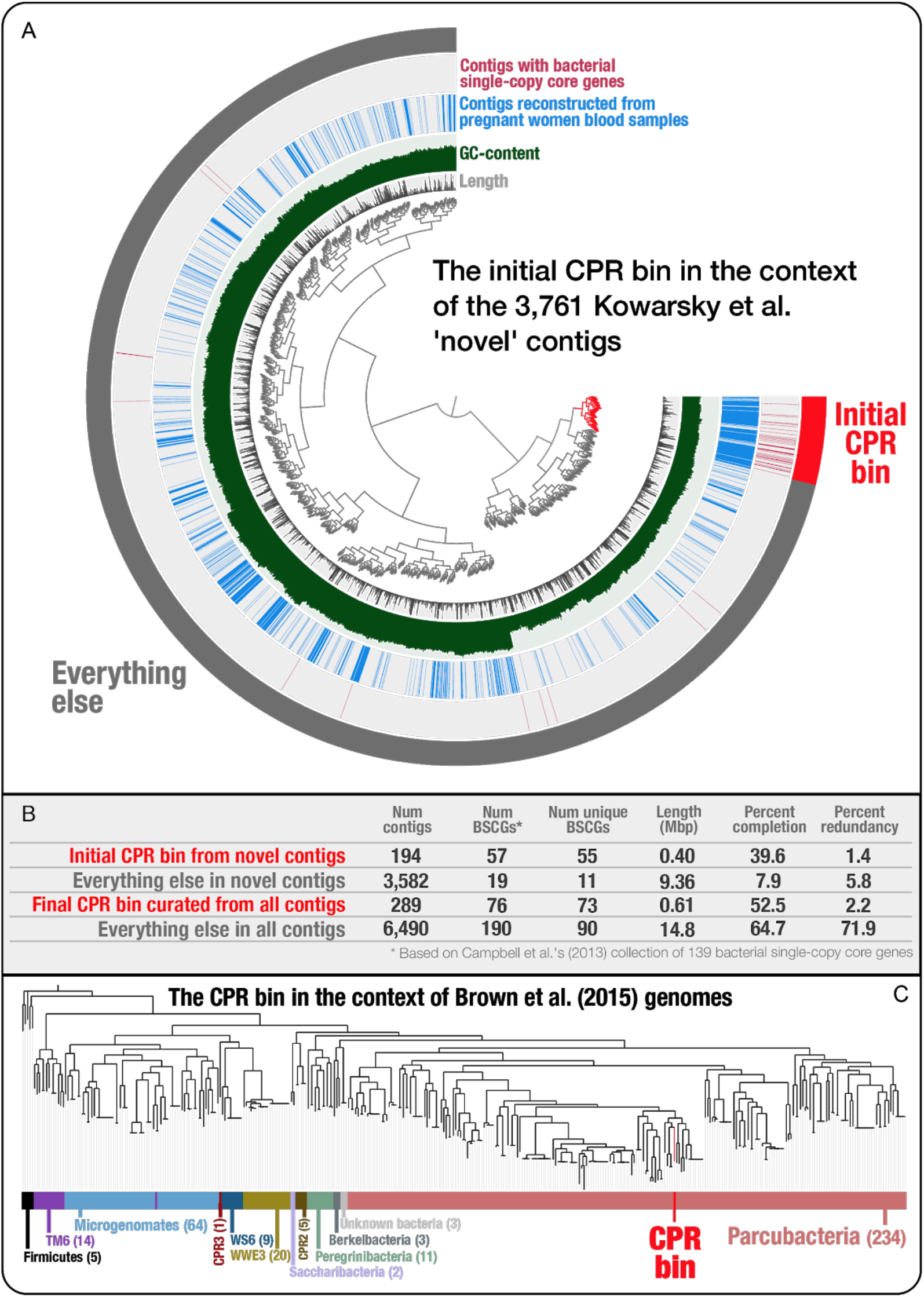
Genome-resolved metagenomics is essential to better predict microbial diversity. (A) The inner dendrogram in Panel A displays the hierarchical clustering of 3,761 ‘novel’ Kowarsky et al. contigs based on their tetranucleotide frequency (using Euclidean distance and Ward clustering). While the two inner layers display the length and GC-content of each contig, the outer two layers mark each contig that originates from the assemblies of pregnant women blood samples, and the ones that contain one or more bacterial single-copy core genes. Panel B compares the initial CPR bin and the remaining contigs in the ‘novel’ set, as well as the final CPR bin and the remaining contigs in the entire assembly (which contains both novel and non-novel contigs). Panel C shows the placement of the CPR bin in the context of CPR genomes released by Brown et al. (2015). See http://merenlab.org/data/parcubacterium-in-hbcfdna/ for more details on this case study.

### Yet, binning can be an important source of error

A real danger is that conclusions from draft MAGs may be incorrect due to misbinning (the wrong assignment of a genome fragment from one organism to another). It is critical to not rely on MAGs with high levels of contamination as these will likely yield misleading evolutionary and ecological insights (Bowers et al. 2017; Shaiber and Eren 2019). Misbinning is especially likely if scaffolds are short (e.g., < 5 kbp), where binning signals can be noisy or unreliable. Thus, for better binning performance, it is helpful to use an assembler that includes a scaffolding step (insertion of Ns in gaps between contigs spanned by paired-end reads), such as IDBA_UD (Peng et al. 2012), or metaSPAdes (Nurk et al. 2017). MAGs can also be screened for short scaffolds with, for example, erroneous rRNA genes, which are often misbinned due to their anomalous coverage (especially if the scaffolds are short and the genes are present in multicopy). Bins may also be contaminated by phage and plasmid genome fragments with coincidentally similar coverage or GC content etc.

Completeness and contamination are often estimated using the inventory of expected SCGs in a MAG. A set of SCGs is selected based on their presence in all bacterial genomes, or at least all genomes within a taxonomic group (identified based on the phylogeny). In a genome without contamination they should be present without redundancy. A widely used tool to assay both completeness and contamination is CheckM (Parks et al. 2015), although other methods are in use (Eren et al. 2015; Anantharaman et al. 2016). It has been noted both in the original study and in subsequent studies that CheckM can generate a false sense of bin accuracy, as demonstrated by combining two partial single-cell genome bins (Parks et al. 2015; Becraft et al. 2017). The absence of multiple copies of SCGs does not preclude the presence of fragments from unrelated organisms that will compromise the biological value of the MAGs. While there are tools for interactive visualization of genome bins in a single sample (Laczny et al. 2015; Raveh-Sadka et al. 2015) or across multiple samples (Eren et al. 2015) that enable manual curation opportunities to identify contamination beyond SCG-based estimates, the scalability of this strategy is limited. For example, recently there have been reports of many thousands, even hundreds of thousands, of draft MAGs from public metagenomic datasets (Parks et al. 2017; Almeida et al. 2019; Nayfach et al. 2019; Pasolli et al. 2019). Such large-scale analyses often rely on simplified procedures, e.g., coverage profile of a single sample for binning, use of a single binning algorithm, completeness/contamination estimates based on SCG inventories. As these genomes are readily adopted by the scientific community for a wide variety of investigations, errors due to misbinning will propagate.

### A case study: SCGs can fail to predict the quality of MAGs

In a recent publication, Pasolli et al. used a single-sample assembly approach combined with automatic binning to generate 345,654 MAGs from the human microbiome, of which 154,723 pass a completion and quality threshold based on SCGs (Pasolli et al. 2019). The authors suggest that the quality of the MAGs they have reconstructed through this pipeline was comparable to the quality of genomes from bacterial isolates or MAGs that were manually curated (Pasolli et al. 2019). However, reconstructing MAGs from single metagenomes and the heavy reliance on SCGs to estimate their quality can yield misleading results.

We examined one of the Pasolli et al. MAGs, ‘HMP_2012__SRS023938__bin.39’ (Pasolli et al. 2019) (hereafter referred to as Pasolli MAG), which resolves to the candidate phylum Saccharibacteria (formerly known as TM7), a poorly understood branch of the Tree of Life that contains members that are common in the human oral cavity (Bor et al. 2019). This MAG, 897,719 bp in length with 57 contigs (N50: 34,012 bp) (Table S2), was recovered by Pasolli et al. from a supragingival plaque sample (experiment accession: SRR060355; sample accession: SRS023938) collected and sequenced by the Human Microbiome Project (HMP) (Turnbaugh et al. 2007). Anvi’o estimated the Pasolli MAG to include 84% of bacterial SCGs with very low redundancy (2.8%), in comparison, CheckM reported 63.39% completeness and 0.85% contamination (Table S2).

The HMP dataset included two additional plaque metagenomes from the same individual, providing an opportunity to investigate the distribution patterns of contigs binned together in this MAG across multiple samples from the same person through metagenomic read recruitment. Organizing contigs based on their sequence composition and differential coverage patterns across three samples revealed two distinct clusters (Figure 3), the smaller one of which contained 11 contigs that added up to a total length of 53.5 kbp (Figure 3, outer circle: orange). While the average mean coverage of contigs in these clusters were relatively comparable in the metagenome from which the MAG was reconstructed (24.6× vs 31.1×), the average coverages differed more dramatically in the other two plaque metagenomes (99.4× vs 20.7× in SRS013723 and 9.7× vs 33.56× in SRS078738), which suggest that the emergence of these two clusters was due to the improved signal for differential coverage with the inclusion of additional samples (Figure 3). A BLAST search on the NCBI’s non-redundant database matched genes found in 10 of 11 contigs in the smaller cluster to genomes of *Veillonella* (belonging to Firmicutes; Table S3), a genus that is common to the human oral cavity (Mark Welch et al. 2014) and includes members that are present in multiple oral sites (Eren et al. 2014). Genes in the remaining contig in the smaller cluster lacked a strong match (contig 000000000028, Table S3), yet best matched to genes in *Selenomonas* genomes instead of Saccharibacteria, suggesting that the smaller cluster represented contamination. As these contaminating contigs did not include any SCGs, their inclusion did not influence SCG-based completeness and contamination estimates. Thus, they remained invisible to the quality assessment. While the contamination in this case will unlikely influence the placement of this particular MAG in the Tree of Life due to the lack of SCGs in it, the contamination does change the functional makeup of the MAG: our annotation of 54 genes in the 11 contaminating contigs using the NCBI’s Clusters of Orthologous Groups (COGs) revealed 30 functions that were absent in the MAG after the removal of the contamination (Table S4). In addition to misleading functional profiles, contamination issues often influence ecological insights. Our read recruitment analysis to characterize the distribution of the Pasolli MAG contigs across all 196 plaque and 217 tongue metagenomes from 131 HMP individuals showed that while this Saccharibacteria population appears to be restricted to plaque samples, contigs that contaminated this MAG recruited reads also from the tongue samples (Figure 3, Table S5).

We did not investigate the quality of the full set of 154,723 MAGs described by Pasolli et al. (Pasolli et al. 2019) or the genomes reported in other studies that relied on similar automated strategies (Almeida et al. 2019; Nayfach et al. 2019). Nevertheless, this example demonstrates that SCGs alone cannot predict the lack of contamination in a given MAG or characterize the extent of contamination in genomic collections (see another example in Figure S2). Overall, it is essential for our community to note that computational analyses that rely heavily on SCGs to assess the quality of MAGs can promote erroneous insights.

**Figure 3.**
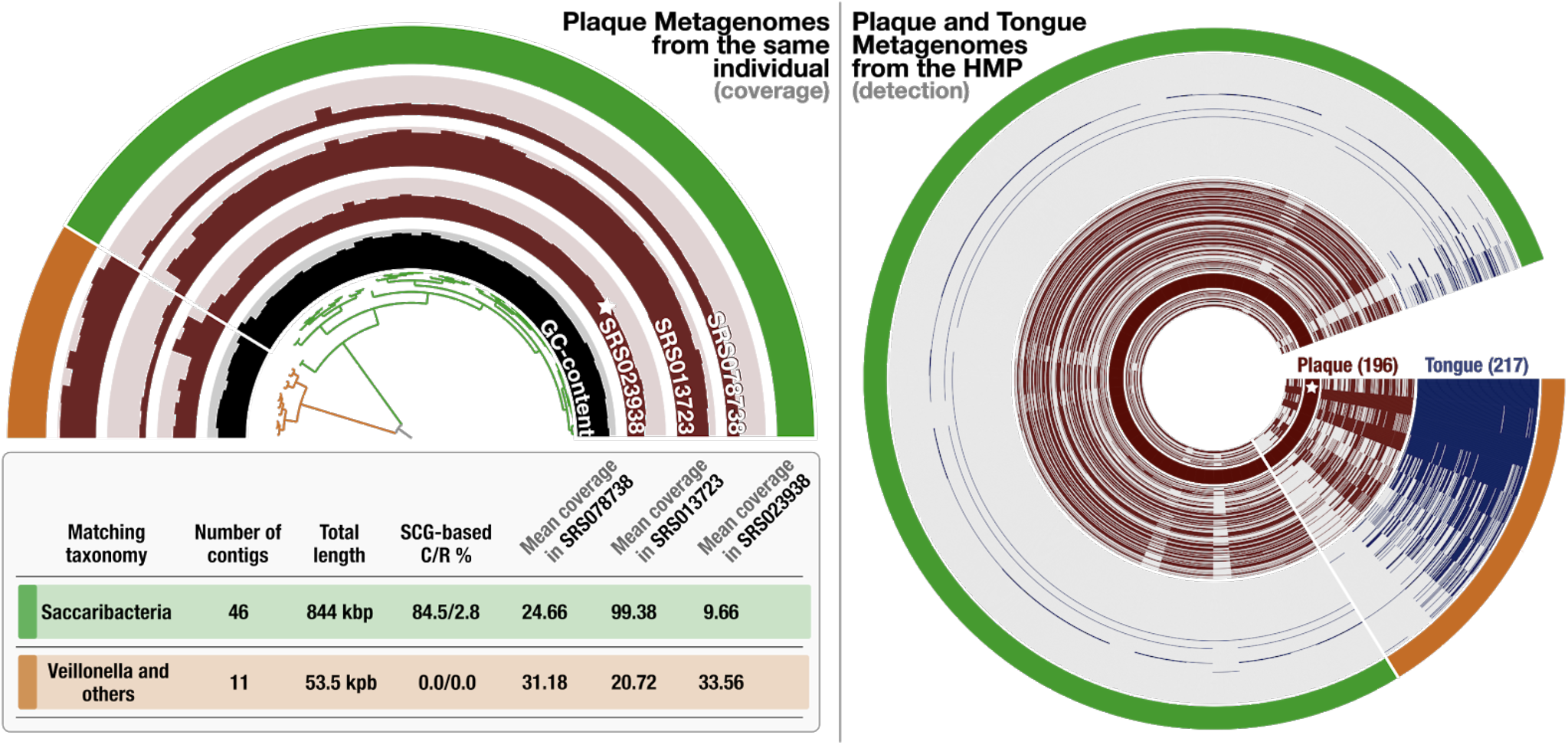
Contamination in MAG without extra copies of SCGs. In the left panel, the half-circle displays the mean coverage of each contig in Pasolli MAG across three plaque metagenomes that belong to the same individual, where the ‘star’ symbol denotes the sample from which the original MAG was reconstructed. The dendrogram in the center represents the hierarchical clustering of the 57 contigs based on their sequence composition and differential mean coverage across the three metagenomes, while the innermost circle displays the GC-content for each contig. The outermost circle marks two clusters: one with 46 contigs (green) and another one with 11 contigs (orange). The table underneath this display summarizes various statistics about these two clusters, including the best matching taxonomy, total length, completion and redundancy (C/R) estimations based on SCGs, and the average mean coverage of each cluster across metagenomes. In the right panel, the distribution of the same contigs and clusters are shown across 196 plaque (brown) and 217 tongue (blue) metagenomes generated by the Human Microbiome Project (HMP). Each concentric circle in this display represents a single metagenome, and data points display the detection of the contigs in Pasolli MAG.

### Genome curation - moving towards complete genomes

The opportunity to recover huge numbers of new genomes from metagenomic datasets motivates the development of new tools to more comprehensively curate draft MAGs, ideally to completion. Although the term ‘complete’ should be reserved for genome sequences with (usually) circular chromosomes reported in single scaffolds, in contemporary genome-resolved metagenomics studies the term is commonly used to describe bacterial and archaeal genomes that have all the expected SCG markers used to evaluate completeness. This use of the term ‘complete’ does not exclude genomes that are extremely fragmented, which can suffer from contamination issues, as we demonstrate above. Here we use the term ‘complete’ explicitly to describe multiple properties of a genome: (1) circular (assuming the chromosome is circular) and single chromosomal sequences, with (2) perfect read coverage support throughout (i.e., the vast majority of bases in mapped reads at any position matches to the consensus base), and (3) no gaps. To avoid any confusion, we will use the term ‘CMAGs’ to describe complete MAGs that meet the three criteria.

One of the first CMAG appeared in 2008, but this was for a bacterium that comprised > 99.9% of the sample (Chivian et al. 2008). A second genome published in the same year was for a candidate phylum bacterium in a bioreactor and was reconstructed by sequencing of a fosmid library (Pelletier et al. 2008). And the third one was a Elusimicrobia genome reconstructed from Termites gut (Hongoh et al. 2008). It was not until 2012 and 2013 that a series of CMAGs from multi-species natural communities began to appear (Iverson et al. 2012; Castelle et al. 2013; Di Rienzi et al. 2013; Kantor et al. 2013). In most cases, these genomes were very close to complete upon *de novo* assembly, although some effort was required to finish them. Near complete *de novo* assembly is a very rare outcome, given that most genomes are assembled using short paired-end reads (e.g., 150 bp with a few hundred base pair insert size). However, given that many samples generate hundreds of draft genomes, very high quality *de novo* assembly of a genome is not uncommon overall. Nevertheless, the curation of even very well assembled MAGs is very rarely undertaken, perhaps due to the involvement of typically manual and generally not well understood steps. Here, we describe the methods that can be used for genome curation and provide examples to illustrate potential caveats along with their likely solutions. Our hope is that the following sections will motivate the development of new tools to enable routine curation of genomes from metagenomes.

### A limited number of published complete metagenome-assembled genomes

To the best of our knowledge, as of 09/10/2019, 59 bacterial and three archaeal CMAGs from microbial community datasets are publicly available (**Table 1**). Of these, four CMAGs were finished using PacBio reads. The published CMAGs are primarily for members of the Candidate Phyla Radiation (CPR; 36 genomes) and DPANN (2 genomes), which have unusually small genomes (average genome size of 1.0 Mbp; Table 1). Other reported CMAGs include those for Proteobacteria (7 genomes), Saganbacteria (WOR-1; 4), Bacteroidetes (two), Candidatus Bipolaricaulota (two), Firmicutes (two), and one from each of Dependentiae (TM6; also small genomes), Elusimicrobia, Melainabacteria, Micrarchaeota, Nitrospirae, Zixibacteria and Candidatus Cloacimonetes (Table 1).

**Table 1.**
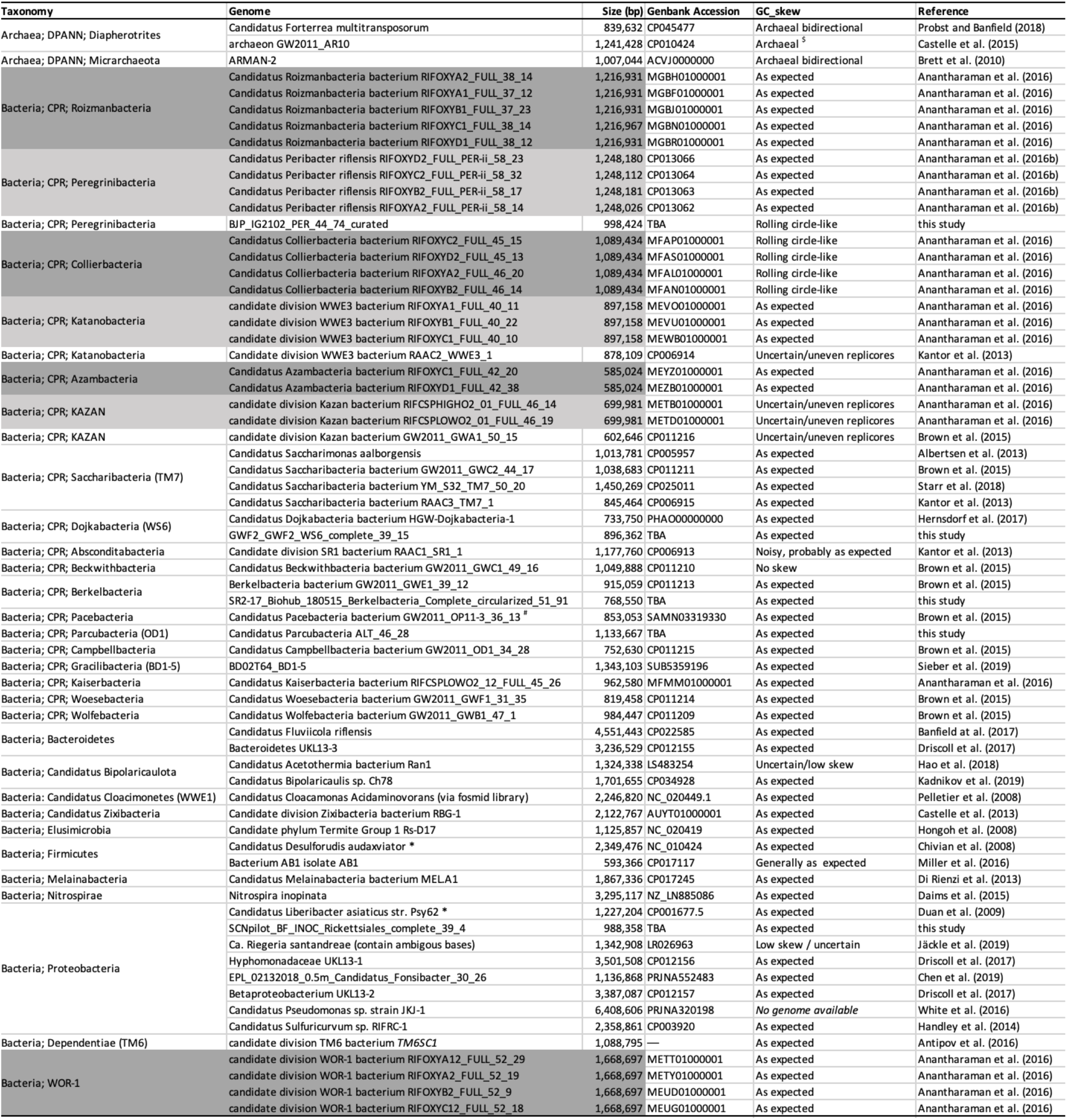
List of complete metagenome-assembled genomes. *The CMAG was reconstructed from a sample with only one organism present. ^#^Wrongly labeled in NCBI as TM6. Note that many genomes exhibit asymmetric patterns of GC skew, which is attributed to uneven length replichores (also seen in isolate genome analysis). ^$^Bidirectional skew patterns are not expected in many archaea. Grey shading indicates essentially identical genomes independently assembled from different samples. To date, CMAGs have been reconstructed for organisms from 30 different phylum-level groups. Five of the listed genomes are complete sequences were completed in the current study.

CMAGs are not limited to bacteria and archaea. Because all of the extracted DNA is sequenced, genomes are also reconstructed for phage and plasmids. In fact, the tool VirSorter (Roux et al. 2015) predicts circularized sequences suitable for verification and curation to remove gaps and local assembly errors. Two recent studies reported unusually large complete phage genomes. In the first case, 15 complete megaphage genomes, each > 540 kbp in length, were reconstructed and curated from human and animal microbiomes (Devoto et al. 2019). In the second case, 35 complete genomes > 200 kbp derived from phage, including the largest phage genomes yet reported (Al-Shayeb et al. 2019). The distinction of these sequences from prophage and the accurate size determinations could not be made without circularized genomes, and the complete, accurate inventory of genes would be precluded with only draft genomes.

### Genome curation: filling scaffolding gaps and removal of local assembly errors

Genome curation requires the identification and correction of local assembly errors and removal of gaps at scaffolding points. However, the exclusion of these steps in current genome-resolved metagenomics studies propagate errors such as incomplete or incorrect protein-coding gene sequences in public databases.

Automatic tools like Gapfiller (Nadalin et al. 2012) may be useful for the filling of ‘N’ gaps at scaffold joins (read pairs should span the gap if the scaffolding was done correctly). Our primary approach to gap filling makes use of unplaced pairs for reads adjacent to the gaps. When reads are mapped to genome fragments that comprise a bin, a file of unplaced paired reads is generated for each fragment. By mapping these unplaced paired reads to the corresponding fragment, it is usually possible to incrementally close the gap (so long as there is sufficient depth of coverage). After the first round of mapping of unplaced paired reads, the consensus sequence must be extended into the gap before remaining unplaced paired reads are remapped. The newly introduced paired reads should be placed at an appropriate distance from their existing pairs, given the fragment insert size. Often a few iterations are needed for gap closure. However, if the gap does not close and no further extension can be accomplished using the existing collection of unplaced pairs, the full metagenomic read dataset can be mapped to the new version of the scaffold and another round of extension performed until the gap is closed.

If a gap cannot be closed using the unplaced paired reads due to low coverage, one solution may be to include reads from another sample in which the same population occurs (this may not be appropriate for some investigations), or by performing a deeper sequencing of the same sample. In other cases, the necessary reads are misplaced, either elsewhere on that scaffold or on another scaffold in the bin. This happens because the reads have been “stolen” thus the true location sequence is not available to be mapped to. This often leads to read pileups with anomalously high frequencies of SNVs in a subset of reads. However, anomalously high read depths can also occur due to mapping of reads from another genome. The misplaced reads can be located based on read names and extracted for gap filling. Other indications of misplacement of reads include read pairs that point outwards (rather than towards each other, as expected) or with unusually long paired read distances. One of these reads is misplaced and the other read normally constrains the region to which the pair must be relocated. Relocation of the misplaced read can often lead to filling of scaffolding gaps. In some cases, gap filling cannot be easily achieved despite sufficient read depth. This can occur, for example, due to complex repeats. Sometimes these repeat regions can be resolved by careful read-by-read analysis, often requiring relocation of reads based on the placement of their pairs as well as sequence identity.

Another important curation step is the removal of local assembly errors (Figure S3). We suspect that these errors are particularly prominent in IDBA_UD assemblies, although it is likely that all assemblers occasionally make local assembly errors. Local assembly errors can be identified because the sequence in that region lacks perfect support, by even one read. The region should be opened up and each read within that region separated to the appropriate side of the new gap (so that all reads match the consensus sequence). Unsupported consensus sequence should be replaced by Ns. The new gap can be filled using the procedure for filling scaffolding gaps, as described above.

A second type of local assembly error is where ‘N’s have been inserted during scaffolding despite overlap between the flanking sequences (Figure S4). We have observed this problem with both IDBA_UD and CLC workbench assemblies. The solution is simply to identify the problem and close the gap, eliminating the Ns and the duplicate sequence.

Another common assembly error involves local repeat regions in which an incorrect number of repeats has been incorporated into the scaffold sequence. This situation may be detected by manual inspection of read mapping profile, as it leads to anomalous read depth over that region. Sometimes the correct number of reads may only be approximated based on the consistency of the coverage within the repeat region and other parts of the scaffold (see example below).

Rarely, in our experience, assemblers create scaffolds that are chimeras of sequences from two different organisms (e.g., (Rojas-Carulla et al. 2019)). These joins typically lack paired read support and/or can be identified by very different coverage values and/or phylogenetic profiles on either side of the join.

Another seemingly rare error involves the artificial concatenation of an identical sequence, sometimes of hundreds of bp in length, repeated up to (or more than) three times. This has been a problem with some sequences of seemingly large phage deposited in public databases (as discussed by Devoto et al. 2019 and Al-Shayeb, Sachdeva et al. 2019). This phenomenon is easily identified by running a repeat finder, a step that should also be included in the curation to completion pipeline (see below).

### From high quality draft sequences to complete genomes from metagenomes

Genome curation to completion is rarely undertaken (Table 1) because there is no single tool available to accomplish it, and there can be confusing complications. The procedure requires the steps described in the prior section as well as extension of scaffolds (or contigs, if no scaffolding step was undertaken) so that they can be joined, ultimately into a single sequence (assuming the genome is a single chromosome). With currently available tools, this is time consuming, sometimes frustrating and often does not result in a CMAG (usually because of indistinguishable multiple options for scaffold joins typically due to repeats such as identical copies of transposons). However, when it can be done, the resulting genome solution should be essentially unique, as we will show below. There is nothing ‘arbitrary’ about the process, except occasionally the choice of which set (usually a pair) of sub-equal locus variants (e.g., SNVs) will represent the final genome. Even in those cases, depending on the availability of multiple appropriate metagenomes for read recruitment analyses, tools for haplotype deconvolution such as DESMAN (Quince et al. 2017) may offer quantitative support for such decisions.

In our experience, the most important first step in the path toward recovery of a CMAG is to start from a well-defined bin that appears to comprise the vast majority of the genome of interest (step 1; Figure 4). As above, this is usually determined based on genome completeness evaluation (step 2; Figure 4) and/or a very strong set of binning signals (see Figure S1 for example). It should be noted that some genomes (e.g., CPR bacteria) may naturally lack certain SCGs that are otherwise considered universal in other bacteria (Brown et al. 2015), and may require a modified list of universal SCGs such as those proposed for CPR genomes for more accurate evaluations of completion (Anantharaman et al. 2016). Importantly, the targeted bin should be polished to remove contamination scaffolds, as noted above (step 3; Figure 4).

**Figure 4.**
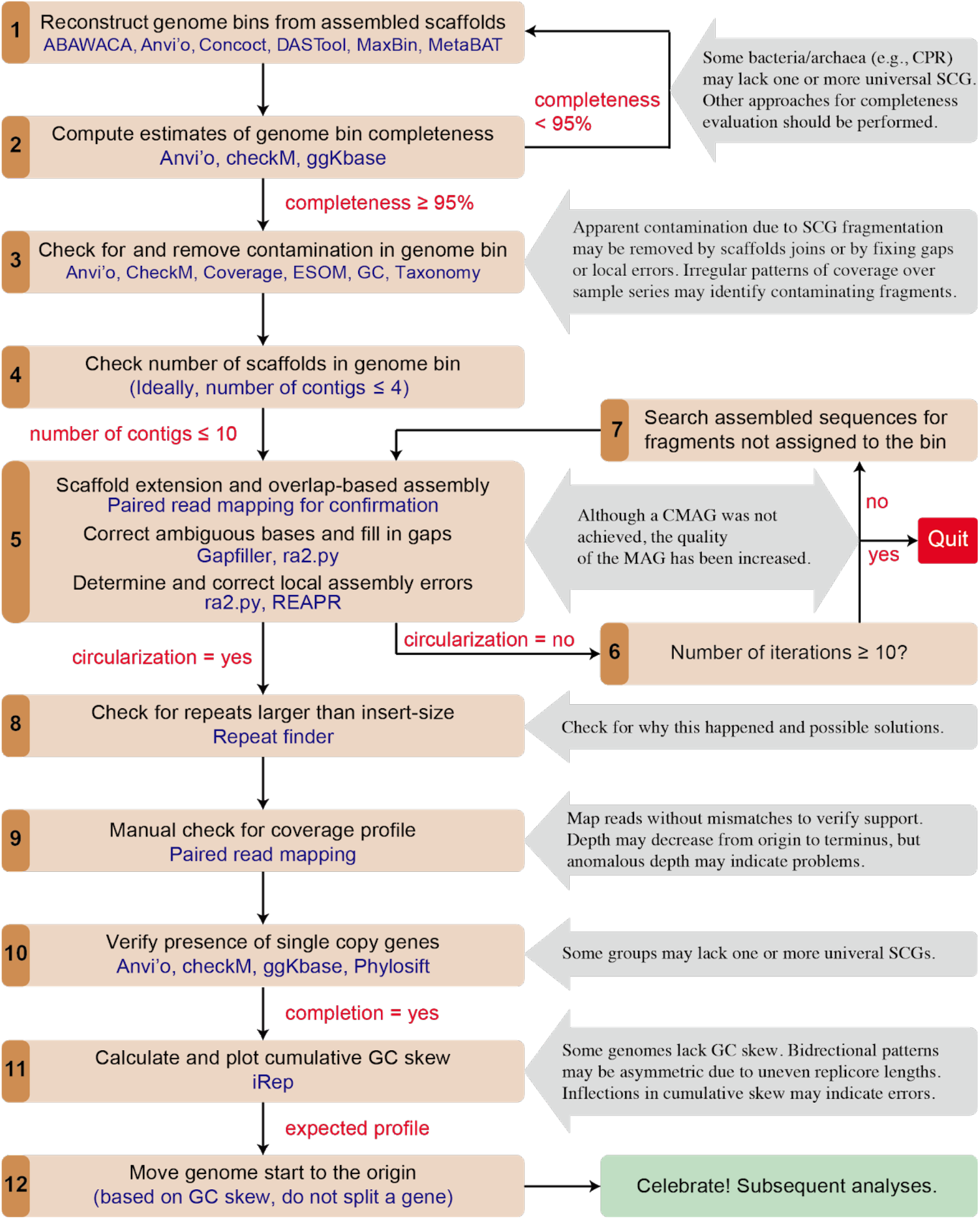
The workflow for generating curated and complete genomes from metagenomes. Steps are shown in black font and the tools or information used in blue font. Notes for procedures are shown in gray boxes.

Given currently available tools, it is probably wise to choose a bin with no more than 10 pieces (step 4; Figure 4), although a MAG with larger number of scaffolds can be curated to completion if necessary (Chen et al. 2019). The best possible case is when the genome is *de novo* assembled into a single piece. In some cases, the genome is already circularized, based on overlap sequences at the scaffold ends, with paired-end reads that span the scaffold ends. Although rare, this does occur, mostly for small genomes (e.g., Saccharibacteria; Albertsen et al. 2013; Starr et al. 2018). In other cases, a modest amount of end extension may be required for circularization (see below). The single scaffold should be checked for complete coverage and support of the consensus. Gaps or local assembly errors must be dealt with before the genome is classified as curated and complete (some additional checks are described below).

Some assemblers (e.g., IDBA_UD, metaSPAdes) retain sequences that are non-unique at scaffold ends. Assembly termination presumably happens because assembly algorithms are designed to stop at points of uncertainty rather than risk making incorrect joins (Figure S5A). Incidentally, as different assemblers can yield different results, there can be value in comparing the results for the same data assembled using different tools and/or parameters (see examples below). Also, in some cases, assembly of scaffolds representing the same organism (or a closely related organism) from a related sample, could help scaffold extension and/or linkage (Figure S5B). Potential scaffold joins can be made by identifying perfect overlaps at the ends of scaffolds of a MAG (“overlap-based assembly”, Step 5; Figure 4). Often, the length of perfect overlap of scaffolds assembled using IDBA_UD and metaSPAdes is n-1 or n, respectively, where n is the largest k-mer size used in *de novo* assembly. Although the assembler chose not to make these joins (possibly due to confusion involving even a single read), seemingly unique joins involving scaffolds in a bin can be made tentatively during curation. Ultimately, non-unique joins can be eliminated or resolved at the end of the curation process. It is important to note that non-uniqueness of a join may not be evident in an initial scaffold set due to failure to include a relevant scaffold in the bin or lack of *de novo* assembly of relevant regions. Thus, it is important to test for repeated regions that cannot be spanned by paired reads at the end of curation (either in the potentially complete genome or curated scaffold set if completion is not achieved). Failure to identify perfect repeats can also lead to problems in isolate genomes, as we show below.

Scaffolds within a bin that do not overlap at the start of curation may be joined after one or more rounds of scaffold extension (Figure S6). This process of extending, joining and remapping may continue until all fragments comprise a single circularized sequence. It should be noted that read by read scaffold extension is very time consuming. If an extended scaffold cannot be joined to another scaffold after a few rounds of extension it may be worth testing for an additional scaffold (possibly small, thus easily missed by binning) by searching the full metagenome for overlaps (steps 6 and 7, Figure 4). Sometimes, the failure of scaffold extension is due to missing paired reads, which may be found at the end of another fragment. If they are pointing out but the sequences cannot be joined based on end overlap, a scaffolding gap can be inserted in the joined sequence (reverse complementing one of the scaffolds may be necessary). Closure of the new scaffolding gap uses the approach described above.

During the attempt to obtain a circularized sequence, it is important to note that if the genome has a single pair of duplicated sequences that are larger than can be spanned by paired reads, a reasonable solution can be found if the genome bin is curated into just two pieces. In this case, the only solutions are either resolution into two chromosomes or generation of a single genome (Figure S7). This observation underlines the importance of curation from a high quality bin, as curation from a full metagenome would leave open the existence of other scaffolds that also bear that repeat.

Once the genome is circularized, it is important to check for repeats larger than spanned by paired-end reads (as noted above; step 8, Figure 4). Assuming a seemingly CMAG is achieved, several steps to further verify the accuracy of the assembly path may be warranted (step 9, Figure 4). First, reads may be mapped to the sequence allowing no mismatches to confirm no coverage gaps due to base miscalls and to verify that no region has abnormal coverage. Tools that provide interactive visualization and inspection of coverage patterns, such as anvi’o, Geneious (Kearse et al. 2012) or ‘Integrative Genomics Viewer’ (Robinson et al. 2011), may be used for this task. Second, we advocate verification of paired read placements over the entire assembly to check for problem areas that may have been missed in automated procedures. Abnormally low coverage may be due to a subpopulation variant, whereas higher than expected coverage could indicate the existence of a block of sequence that was pinched out from the genome at a repeated region. Systematic decline of coverage from origin to terminus of replication is expected if genome replication was ongoing at the time of sampling (see below). Third, the presence of expected genes (e.g., universal SCGs) should be verified. The genome can be classified using phylogenetic analyses (e.g., based on 16S rRNA gene or concatenated ribosomal proteins sequences; step 10, Figure 4). After the completion of MAG, the start of the genome should be moved to the non-coding region near the origin of replication (steps 11 and 12, Figure 4). See below for details regarding how GC skew can be used to locate the origin.

An important consideration in genome curation to completion is knowing when to give up. In some cases, failure to circularize after a few rounds of curation may be an indication that the effort could be better invested in other activities. If alternative assembly paths that cannot be distinguished by the unique placement of paired reads are identified, failure may be on the horizon. However, as noted above (Figure S7), it can be appropriate to continue curation as a final unique solution may be possible even in the presence of a repeat that cannot be spanned by paired reads.

### Case studies illustrating the curation of draft MAGs

Here we illustrate how a draft MAG can be curated to completion or into better quality status, with step-by-step procedures detailed in the Supplementary Information.

#### Case one, curation of a CPR genome to completion

ALT_04162018_0_2um_scaffold_13, length of 1,128,909 bp, was the only scaffold in the binned MAG (i.e., bin.56) from MetaBAT (Figure S8). CheckM reported 70.1% completeness without contamination, and preliminary analyses based on 16S rRNA and rpS3 genes identified it as a Parcubacteria genome. This genome was likely near complete based on the detection of all CPR universal SCGs, though we did not identify overlap at the ends of the scaffold that would circularize it. This scaffold could be circularized after a single round of scaffold end extension, with read pairs placed at the ends of the scaffold. In fact, we found two very small assembled sequences that were variants of each other and both could be used for circularization. The non-uniqueness of this region terminated the original assembly. We chose the dominant variant to represent the population genome. No repeat sequence longer than the sequencing insert size was detected. A total of 13 local assembly errors were reported by ra2.py. All these errors were manually fixed and validated, including a complicated error in the sequence of a protein-coding gene that contains multiple repeat regions. The complete genome has a length of 1,133,667 bp, and encodes 1,147 protein-coding genes, 47 tRNA and a copy each of 5S/16S/23S rRNA genes.

#### Case two, curation of a Betaproteobacteria genome without completion

Bin.19 contained seven scaffolds (3.6 Mbp in size), and was evaluated by CheckM to be 98.42% complete with 0.12% contamination (Figure S9). Analyses of the 16S rRNA gene sequence indicated it was a Betaproteobacteria (92% similarity to that of *Sulfuricella denitrificans* skB26). After the first round of scaffold extension and assembly, only two scaffolds could be combined (i.e., scaffold 21 and 25). We searched for the pieces that could be used to link the scaffolds together using the newly extended parts of the scaffolds via BLASTn against the whole scaffold set. This approach retrieved four short (584-1191 bp in length) and one longer piece (15,678 bp in length) that encodes several bacterial universal SCGs including rpS7, rpS12, rpL7/L12, rpL10, rpL1 and rpL11 (which were absent from bin.19), and whose two ends both encode elongation factor Tu (EF-Tu). Two of the four short pieces could be perfectly joined in two possible places to the original scaffold set. Based on comparison with the *Sulfuricella denitrificans* skB26 genome, we hypothesized the linkage patterns for these fragments and then considered the two choices for how the resulting two large genome fragments could be arrayed. The linkage choices were supported based on the overall pattern of GC skew (see below and Figure S9). Technically, however, the bin remains as two contigs with two internal joins unsupported by unique paired read placement. Based on the GC skew of the pair of contigs linked by Ns, the genome is near complete. After the fixation of local assembly errors, it has a total length of 3.72 Mbp, encodes 3,544 protein-coding genes, 41 tRNA and one copy of each of the 5S/16S/23S rRNA genes, and is clearly of higher quality than the original bin due to scaffold extension and correction of local assembly errors.

#### Case three, curation of a published incomplete genome to completion

Here we completed a published curated (for local assembly errors) but incomplete genome belonging to the order *Rickettsiales* (Kantor et al. 2017). This genome was assembled *de novo* into a single circularizable 988 kbp scaffold, with two closely spaced gaps (Figure S10a). Closing of these gaps required relocation of unplaced paired reads (Figures S10b and c).

In addition to the above-mentioned case studies, we curated three additional bacterial genomes to completion as part of our methods refinement. These genomes are listed in **Table 1**.

### Using GC skew as a metric for checking genome correctness

GC skew is a form of compositional bias (imbalance of guanosine (G) relative to cytosine (C) on a DNA strand) that is an inherent feature of many microbial genomes, although some are known to display little or no GC skew (e.g., certain Cyanobacteria, (Nakamura 2002)). The phenomenon of strand-specific composition was described by Lobry (Lobry 1996), who observed that the relative GC skew changes the sign crossing the *oriC* and *terC* regions. Thus, the inflection point in genome GC skew at the origin of replication is often close to the dnaA gene and typically contains a small repeat array. GC skew is calculated as (G−C/G+C) for a sliding window along the entire length of the genome (suggested window=1000 bp, slide=10 bp). The skew is also often summed along the sequence to calculate cumulative GC skew. This was proposed by Grigoriev (Grigoriev 1998), who showed that the calculation of the cumulative GC skew over sequential windows is an effective way to visualize the location of the origin and terminus of replication. For complete genomes, the GC skew is often presented starting at the origin of replication, proceeding through the terminus and back to the origin (i.e., as if the chromosome was linear). The pattern of the cumulative GC skew, where the function peaks at the terminus of replication, indicates that the genome undergoes bidirectional replication. The pattern is fairly symmetrical unless the replichores are of uneven lengths. Because the magnitude of the cumulative GC skew varies from genome to genome, the magnitude of the skew could potentially be used as a binning signal.

The explanation for the origin of GC skew is not fully agreed upon. It may arise in large part due to differential mutation rates on the leading and lagging strands of DNA. Enrichment in G over C occurs due to C deamination to thymine (C->T), the rates of which can increase at least 100-fold when the DNA is in a single stranded state. In the process of DNA replication, the leading strand remains single stranded while the paired bases are incorporated by the DNA polymerase into its complementary strand. However, the Okazaki fragments on the lagging strand protect a fraction of the DNA from deamination. Thus, the leading strand becomes enriched in G relative to C compared to the lagging strand. The magnitude of the GC skew can be impacted by the speed of the DNA polymerase processivity (which impacts the length of time that the DNA is single stranded) and the length of the Okazaki fragments. GC skew has been linked to strand coding bias (Rocha et al. 1999). Concentration of genes on the leading strand would afford protection against non-synonymous mutations (as C->T mutations in the wobble position of codons are always synonymous), whereas G->A on the lagging strand (following C->T on the leading strand) in two cases results in nonsynonymous mutations (AUA for Ile vs. AUG for Met, and UGA for stop codon vs. UGG for Trp). The potential for deamination in the non-coding strand during transcription, another source of GC skew, would also favor genes on the leading strand. GC skew persists because the leading strand is maintained as such through subsequent replication events.

Given that a well-defined pattern of GC skew is anticipated across many bacterial (and some archaeal) genomes, we wondered whether plots of cumulative GC skew for putative complete genomes can be confidently used to test for genome assembly errors. For this metric to be useful, it would be imperative to establish the extent to which GC skew is indeed a feature of complete bacterial genomes. To our knowledge, the now extensive set of complete isolate genomes has not been leveraged to do this.

We undertook benchmarking of GC skew, and more specifically cumulative GC skew, using all ~7000 complete genomes in the RefSeq database. We found that the majority of RefSeq bacterial genomes show the expected pattern of cumulative GC skew. Interestingly, the magnitude of the origin to terminus skew varies substantially, from ± 0.4 excess G relative to C to close to zero (Figure S11). A small subset of the ~7000 complete genomes essentially lack GC skew (as reported for some Cyanobacteria, see above) (Table S6). Poorly defined (noisy) patterns are often associated with low total cumulative skew. About 15% of genomes have notably asymmetric patterns (i.e., the cumulative skew is substantially larger for one half of the chromosome relative to the other), presumably because the two replichores are of substantially uneven length. Moreover, some bacterial genomes had a GC skew pattern indicating rolling circle replication (Table S7). Interestingly, we did not detect a strong correlation between the magnitude of GC skew and bias for genes on the leading strand.

Some complete genomes have quite aberrant skew patterns, with inversions in the cumulative skew within a single replichore or exceedingly uneven predicted replichore lengths. We considered the possibility that a subset of these isolate genomes may contain mis-assemblies. Such a phenomenon was already shown by Olm et al. in the case of a *Citrobacter koseri* isolate genome that was clearly wrongly assembled across rRNA operons (and a PacBio assembly for a closely related strain showed the expected pattern of cumulative GC skew) (Olm et al. 2017). To test for the possibility that these other complete genomes contained errors, we posited that mis-assemblies would likely occur at perfect repeats that are longer than the distance spanned by paired reads. Further, we predicted that the pair of repeats flanking the wrongly assembled sequence region would be in reverse complement orientations so that the intervening DNA segment could be flipped at the repeats and that the flipped version would exhibit the expected GC skew pattern. In five of twelve cases that we scrutinized it was possible to show that reverse complementing the sequence spanned by repeats indeed resulted in genomes with exactly the expected form of cumulative GC skew (Figures 5, S12 and S13). In one case, i.e., *Flavobacterium johnsoniae* UW101 (NC_009441.1), the original assembly notes indicated assembly uncertainty (although the complete genome was deposited at NCBI).

We acknowledge the possibility that a recent major rearrangement could also give rise to inflexions in GC skew, however major rearrangements typically have a well-defined placement relative to the origin of replication that is inconsistent with the patterns observed (Eisen et al. 2000). Although we cannot state that these isolate genomes are wrongly assembled, we suggest that it is a distinct possibility. Incorrect assemblies in isolate genomes can be of high significance, given the trust placed in them for evolutionary and metabolic analyses that make use of synteny and gene context. They are also used as references for calculation of growth rates via the PTR method (Korem et al. 2015), and incorrect reference sequences will corrupt such measurements.

**Figure 5.**
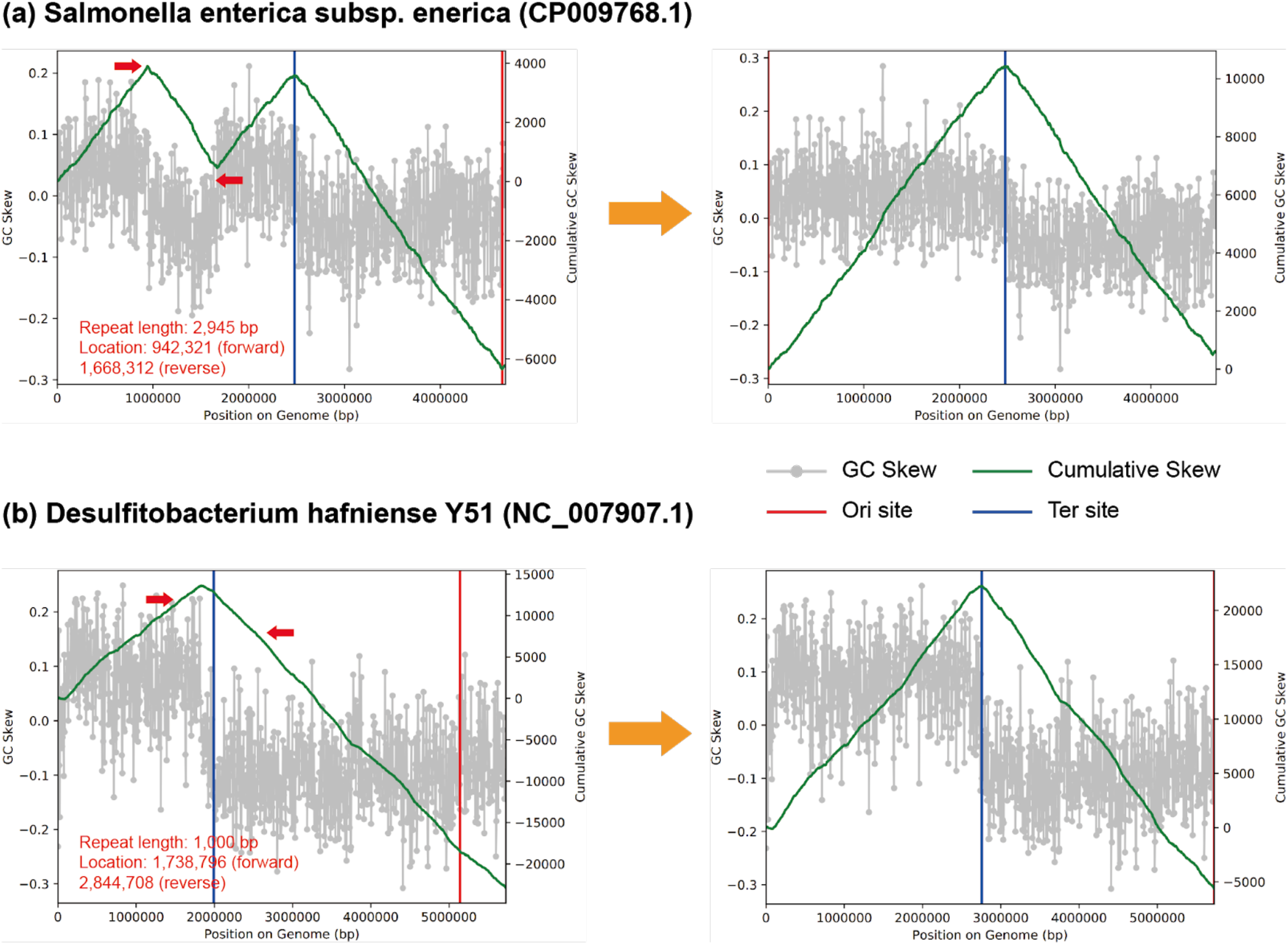
Examples of probable assembly errors in RefSeq bacterial genomes. The diagrams show the GC skew (grey) and cumulative GC skew (green line) of the original (left) and the modified (right) versions of the genomes (all calculated with window size of 1000 bp, and slide size of 10 bp). The location and direction of repeat sequences leading to the abnormal GC skew are indicated by red arrows. After flipping the repeat-bounded sequences the genomes show the pattern expected for genomes that undergo bidirectional replication (right). For more examples, see Figures S12 and S13.

It is well known that some archaea replicate their genomes from multiple origins (Barry and Bell 2006). In such cases, the cumulative GC skew pattern is not a useful test of overall genome accuracy. However, some archaea do show the peaked pattern that is typical of bacteria, thus indicative of bidirectional replication. Overall, we found 18 of 224 RefSeq archaeal genomes tested that show this pattern, and all of them are Euryarchaeota (Table S8). In addition, this pattern was reported for a DPANN archaeon (Probst and Banfield 2018).

### Other approaches, future opportunities and challenges

#### Single-cell genomics

Microbial single-cell sequencing is a family of strategies that typically use microfluidics and whole-genome amplification to physically isolate individual cells and sequence their genomes without cultivation (Stepanauskas 2012). The resulting single-amplified genomes (SAGs) can offer critical insights into microbial lifestyles (Swan et al. 2011), and shed light on intra-population structures of complex microbial consortia (Kashtan et al. 2014) or naturally occurring host-virus interactions (Labonté et al. 2015), where short-read and assembly-based strategies may not be effective. However, state-of-the-art single-cell sequencing strategies typically generate highly fragmented and incomplete genomes due to the need for random amplification arising from small quantities of DNA present in a single cell (Kalisky and Quake 2011). In some cases, sequences from other organisms may contaminate individual wells (Rinke et al. 2013), in other cases combining sequences from different cells into single draft genomes based on sequence identity thresholds of phylogenetic markers (i.e. > 97% 16S rRNA identity; (Rinke et al. 2013)), may result in hybrid genomes. In fact these hybrids are potentially from different species, given that many consider 97.9% 16S rRNA sequence divergence as a proxy for the species boundary (Newton et al. 2007; Garcia et al. 2018). Interestingly, Probst et al. indicate that although the cells are often chosen for single cell sequencing based on their amplified 16S rRNA genes, the sequences recovered do not always match the amplified genes (Probst et al. 2018). Some of these problems may be ameliorated with additional steps of binning and refinement, and similar to MAGs, SAGs can also be curated to completion as demonstrated by at least one study that used long (Sanger) reads in conjunction with short read assemblies (Woyke et al. 2010). Given the fast pace of improvements in microfluidics technologies as well as whole-genome amplification and sequencing chemistry (Woyke et al. 2017), we anticipate that single-cell genomics will continue to gain popularity and its joint use with other genome-resolved metagenomics strategies will become increasingly frequent.

#### Complete genomes from long-reads

Among the published CMAGs, four were obtained by assembly of PacBio reads, including three proteobacterial and one Bacteroidetes genome (White et al. 2016; Driscoll et al. 2017). Especially Oxford Nanopore Technologies offers affordable, easy-to-operate, and portable sequencers for long-read sequencing. While improving, errors from nanopore sequencing can dramatically exceed state-of-the-art short read sequencing (Laver et al. 2015), however, new approaches for long-read correction (Rang et al. 2018; Arumugam et al. 2019), hybrid assembly (Wick et al. 2017), and mock community standards (Nicholls et al. 2019) are emerging. Short read-based assembly strategies often report fragmented contigs due to repeat elements that exceed short read lengths, which is an issue long-read sequencing overcomes, improving the quality of genomes from metagenomics (Arumugam et al. 2019). We anticipate that the combination of short reads and long-reads sequencing will be an increasingly common strategy for recovery of highly curated and complete genomes from microbial community samples.

#### Chromosome conformation capture method

The chromosome conformation capture (i.e., 3C) is a method that enables the determination of physical contacts between different regions of a chromosome and between the different chromosomes of a cell (Dekker et al. 2002). The initial applications of this strategy focused on eukaryotic genomes and revealed, for example, the folding principles (Lieberman-Aiden et al. 2009) and the chromatin looping (Rao et al. 2014) of the human genome. The 3C approach has recently been developed into multiple derivative proximity ligation methods such as Hi-C (Lieberman-Aiden et al. 2009) and meta3C (Marbouty et al. 2014), and applied to individual microbial populations (Le et al. 2013) as well as complex assemblages of environmental microbes (Marbouty et al. 2014). As these approaches offer physical linkage between DNA fragments that are proximal to each other, they can improve metagenomic binning (Baudry et al. 2019; DeMaere and Darling 2019). While promising, the additional complexity of library preparations and additional cost due to the need for separate metagenomic libraries (Liu and Darling 2015) prevent their routine application to metagenomic studies. In addition, distinct populations that are in close proximity in the input sample and repeat sequences may yield misleading contact signals and result in chimeric assemblies (Marbouty and Koszul 2015). Nevertheless, the application of proximity ligation strategies to naturally occurring complex microbial consortia can provide important insights (Bickhart et al. 2019; Stalder et al. 2019).

#### Eukaryotes and even macroorganisms

The assembly of draft eukaryotic genomes from shotgun metagenomes is possible, despite the large genome sizes of most eukaryotes. However, eukaryotic MAGs can be readily contaminated by fragments of genomes from coexisting bacteria and archaea (Boothby et al. 2015; Arakawa 2016), so careful evaluation is needed to avoid misleading conclusions (Delmont and Eren 2016). We have found that phylogenetic profiling of contigs based on best matches in reference databases can be an effective way to identify contaminating bacterial and archaeal sequences.

An important step for recovery of reasonable quality eukaryotic genomes from metagenomes is to separate assembled eukaryotic from prokaryotic genome fragments prior to binning. Then, eukaryote-specific gene predictions can be established and gene annotations used to estimate genome completeness. The K-mer-based classifier, EukRep, was developed to accomplish this separation (West et al. 2018). Although eukaryote genome recovery from metagenomes is increasingly reported (Quandt et al. 2015; Mosier et al. 2016; Olm et al. 2019), to our knowledge, none have been extensively curated or completed.

#### High fragmentation of metagenomic scaffolds

A major limitation on the quality of MAGs relates to genome fragmentation. Fragmentation is doubly problematic because small fragments are hard to bin accurately and gaps result in incomplete gene inventories. Fragmentation can arise due to the presence of duplicated sequences (e.g., transposases, rRNA operons), but the most pronounced problems usually are the result of coexisting closely related strains that confuse de Bruijn graph based assemblers (Olson et al. 2017). For example, although *Prochlorococcus* and SAR11 are among the most abundant bacteria in ocean habitats, the co-occurence of closely related strains (Giovannoni 2017) leads to very fragmented MAGs and poor representation in the final datasets (Delmont et al. 2018; Tully et al. 2018). Of the three commonly used metagenomic assemblers, IDBA_UD, MEGAHIT, metaSPAdes (Greenwald et al. 2017), metaSPAdes was best designed to handle micro-variations between fragments from related strains to generate longer composite sequences (Olson et al. 2017). However, care should be taken when undertaking detailed analyses (e.g., biochemical testing) of open reading frames generated in this way as they may be chimeric.

Practically, another approach that can sometimes address the problem of assembly fragmentation due to strain variety is collections of sequences from related samples (e.g., along a geochemical gradient) to identify communities in which there is much reduced complexity of related strains. For example, opportunities can arise due to the recent proliferation of one strain over the background of numerous closely related strains following changes in environmental conditions. In other words, if a genome cannot be recovered from one sample, look for it in related samples. We anticipate that this approach will be most effective for genome recovery from soil environments, where strain diversity can be extreme and environmental heterogeneity provides access to different strain mixtures.

## Conclusions

Genomes derived from metagenomes have advanced our understanding of microbial diversity (Hug et al. 2016; Anantharaman et al. 2016; Parks et al. 2017) and metabolism (e.g., (van Kessel et al. 2015; Anantharaman et al. 2018). However, these genomes are readily adopted by the scientific community for a wide variety of investigations, and errors will propagate. In fact, the proposal of a new nomenclature for large swaths of the tree of life based largely on MAGs (Parks et al. 2018) brings a potential crisis into focus. We conclude that it is imperative that complete, curated genomes are recovered for all major lineages (including those that lack any isolated representative). The increased span of phylogenetic coverage by complete genomes will provide a valuable reference set against which newly recovered genomes can be confidently compared and augment what has been achieved by the isolate-based Genome Encyclopedia of Bacteria and Archaea program (Wu et al. 2009). New complete sequences from previously genomically undescribed lineages will also improve understanding of how protein families and functions are distributed, facilitate more powerful analyses of evolutionary processes such as lateral gene transfer and enable more accurate phylogenetic representations of life’s diversity. Finally, we advocate for the development of methods to routinely curate assemblies and draft genomes (if not to completion) at scale to ensure the accuracy of evolutionary and ecosystem insights.

## Methods

### Preparation of MAGs as examples for genome curation

This study includes two MAGs were not previously published, as examples for genome curation. These genomes were assembled from samples collected in a mine tailings impoundment (Manitoba, Canada). The raw reads of metagenomic sequencing were filtered to remove Illumina adapters, PhiX and other Illumina trace contaminants with BBTools (Bushnell 2018), and low-quality bases and reads using Sickle (version 1.33; https://github.com/najoshi/sickle). The high-quality reads were assembled using both IDBA_UD (Peng et al. 2012) and metaSPades (Nurk et al. 2017). For a given sample, the quality trimmed reads were mapped to the assembled scaffolds using bowtie2 with default parameters (Langmead and Salzberg 2012). The coverage of each scaffold was calculated as the total number of bases mapped to it divided by its length. The protein-coding genes were predicted from the scaffolds using Prodigal (Hyatt et al. 2010), and searched against KEGG, UniRef100 and UniProt for annotation. The 16S rRNA gene was predicted using a HMM model, as previously described (Brown et al. 2015). The tRNAs were predicted using tRNAscan-SE 2.0 (Lowe and Chan 2016). For each sample, scaffolds with a minimum length of 2.5 kbp were assigned to preliminary draft genome bins using MetaBAT with default parameters (Kang et al. 2015), with both tetranucleotide frequencies (TNF) and coverage profile of scaffolds (from multiple samples) considered. The scaffolds from the obtained bins and the unbinned scaffolds with a minimum length of 1 kbp were uploaded to ggKbase (http://ggkbase.berkeley.edu/). The genome bins were evaluated based on the consistency of GC content, coverage and taxonomic information and scaffolds identified as contaminants were removed.

### GC skew evaluation of RefSeq genomes

We analyzed all the NCBI RefSeq genomes downloaded on May 10th, 2017 for GC Skew. Both skew and cumulative skew were calculated and patterns displayed using the publicly available program gc_skew.py (https://github.com/christophertbrown/iRep) (Brown et al. 2016).

### Refinement of the CPR genome from blood

For initial characterization of the CPR bin we used the contigs made publicly available as the ‘Dataset S6’ in the original study (Kowarsky et al. 2017). These contigs represent what remained after the removal of contigs with matches to sequences in any existing public databases (Kowarsky et al. 2017); we will refer to these contigs as ‘novel contigs’. In our study we also had access to the remaining contigs, and we will refer to this dataset as ‘all contigs’.

For binning and refinement of the CPR genome, and metagenomic read recruitment analyses, we used anvi’o v5.5 to generate a contigs database from the novel contigs using the program ‘anvi-gen-contigs-database’, which recovered the tetranucleotide frequencies for each contig, used Prodigal v2.6.3 (Hyatt et al. 2010) with default settings to identify open reading frames, and HMMER v3.2.1 (Eddy 2011) to identify matching genes in our contigs to bacterial single-copy core genes by (Campbell et al. 2013). To visualize all novel contigs we used the program ‘anvi-interactive’, which computed a hierarchical clustering dendrogram for contigs using Euclidean distance and Ward linkage based on their tetranucleotide frequency (TNF), and displayed additional data layers of contig cohort origin and HMM hits we supplied to the program as a TAB-delimited additional data file. We manually selected a branch of contigs that created a coherent cluster based on the TNF data and the occurrence of bacterial single-copy core genes. While this procedure allowed us to identify an initial genome bin with modest completion, it’s comprehensiveness and purity was questionable since our binning effort (1) utilized only the novel contigs from the (Kowarsky et al. 2017), which were a subset of all contigs assembled, and (2) only employed tetranucleotide signatures to identify the genome bin, which can introduce contamination as the sequence signatures of short fragments of DNA can be noisy. To address these issues, we first acquired the remaining 3,002 contigs that were not included in the original study (Kowarsky et al. 2017) and that might be derived from the same blood-associated CPR population. Then, we used all blood metagenomes for a read recruitment analysis. This analysis allowed us to identify contigs from the non-novel contig collection that match to the distribution patterns of the initial CPR bin. Since the coverage of this population was extremely low, we used a special clustering configuration for anvi’o to use ‘differential detection’ rather than ‘differential coverage’ (see the reproducible workflow for details). This analysis resulted in contigs with similar detection patterns across all metagenomes. We summarized this final collection of contigs using ‘anvi-summarize’, which gave access to the FASTA file for the bin. Anvi’o automated workflows (http://merenlab.org/2018/07/09/anvio-snakemake-workflows/) that use snakemake (Köster and Rahmann 2012) performed all read recruitment analyses with Bowtie v2.3.4 (Langmead and Salzberg 2012). We profiled all mapping results using anvi’o following the analysis steps outlined in Eren et al. (Eren et al. 2015).

To put our CPR bin into the phylogenetic context of the other available CPR genomes, we used the 797 metagenome-assembled CPR genomes (Brown et al. 2015). We used the anvi’o program ‘anvi-get-sequences-for-hmm-hits’ to (1) collect the 21 amino acid sequences found in the CPR bin (Ribosomal_L10, Ribosomal_L11, Ribosomal_L11_N, Ribosomal_L13, Ribosomal_L14, Ribosomal_L17, Ribosomal_L20, Ribosomal_L21p, Ribosomal_L27, Ribosomal_L32p, Ribosomal_L5_C, Ribosomal_L9_C, Ribosomal_L9_N, Ribosomal_S11, Ribosomal_S13, Ribosomal_S16, Ribosomal_S2, Ribosomal_S20p, Ribosomal_S4, Ribosomal_S7, Ribosomal_S9) from all genomes, (2) align them individually, (3) concatenate genes that belong to the same genome, and (4) report them as a FASTA file. Some of the key parameters we used with this program included ‘--hmm-source Campbell_et_al’ to use the single-copy core gene collection defined by Campbell et al. (Campbell et al. 2013), ‘--align-with famsa’ to use FAMSA (Deorowicz et al. 2016) to align sequences for each ribosomal protein, ‘--return-best-hit’ to get only the most significant HMM hit if a given ribosomal protein found in multiple copies in a given genome, and ‘--max-num-genes-missing-from-bin 3’ to omit genomes that miss more than 3 of the 21 genes listed. We used trimAl v1.4.rev22 (Capella-Gutiérrez et al. 2009) to remove positions that were gaps in more than 50% of the genes in the alignment (-gt 0.50), IQ-TREE v1.5.5 (Nguyen et al. 2015) with the ‘WAG’ general matrix model (Whelan and Goldman 2001) to infer the maximum likelihood tree, and anvi’o to visualize the output.

### Refinement of the Pasolli MAG

We downloaded the Pasolli MAG (‘HMP_2012__SRS023938__bin_39’) from http://opendata.lifebit.ai/table/?project=SGB and the 481 HMP oral metagenomes from the HMP FTP server (ftp://public-ftp.hmpdacc.org/Illumina/). We used anvi’o v6 and the Snakemake-based (Köster and Rahmann 2012) program ‘anvi-run-workflow’ to run the anvi’o metagenomics workflow (Eren et al. 2015). Briefly, we generated a contigs database from the Pasolli MAG FASTA file by running ‘anvi-gen-contigs-database’, during which anvi’o calculates tetra-nucleotide frequencies for each contig, and Prodigal (Hyatt et al. 2010) to identify genes. In order to estimate the completion and redundancy of the Pasolli MAG based on SCGs, we used the program ‘anvi-run-hmms’ with the default HMM profiles, which include 71 bacterial SCGs (HMMs described in anvi’o v6), and annotated genes with functions using ‘anvi-run-ncbi-cogs’ which searches amino-acid sequences using blastp v2.7.1+ (Altschul et al. 1990) against the December 2014 release of the COG database (Tatusov et al. 2000). We mapped the paired-end reads from the 481 HMP metagenomes to the Pasolli MAG using bowtie v2.2.6 with default parameters (Langmead and Salzberg 2012) and converted the mapping output to BAM files using samtools v1.9 (Li et al. 2009). We used ‘anvi-profile’ to generate profile databases from BAM files, in which coverage and detection statistics for contigs in each metagenome were stored. We used ‘anvi-merge’ to merge the anvi’o profile databases of (1) only the 3 plaque metagenomes of HMP individual 159268001, which includes the sample from which the Pasolli MAG was constructed (sample accession SRS023938), and (2) all 481 HMP oral metagenomes. In order to manually refine the Pasolli MAG, we ran the anvi’o interactive interface using ‘anvi-interactive’ with the merged anvi’o profile database that included only the three plaque metagenomes of HMP individual 159268001. Refinement was done using hierarchical clustering of the contigs based on sequence composition and differential coverage using Euclidean distance and Ward’s method. To estimate the taxonomic assignment we blasted the protein sequences of genes in the 11 contigs identified as contamination against the NCBI’s non-redundant protein sequences database. To visualize the detection values of the contigs of the Pasolli MAG across all 481 HMP oral metagenomes we used the full merged profile database and the program ‘anvi-interactive’. We used ‘anvi-summarize’ to generate tabular summaries of detection and coverage information of the refined Saccharibacteira bin and the 11 contigs of contamination across the 481 metagenomes.

### Data access

All sequencing data described in this manuscript is available at the NCBI Genbank under accession numbers provided in Table 1.

## Acknowledgements

We thank Brian C. Thomas, Matthew R. Olm, Christopher T. Brown, Alla Lapidus and Christian M.K. Sieber for helpful discussions and Steven Quake and Mark Kowarsky for providing access to unreleased sequences from their cell-free blood study. This work was supported by the Genome Canada Large Scale Applied Research Program and Ontario Research Fund – Research Excellence grants to L.A.W., Lawrence Berkeley National Laboratory’s Watershed Function Scientific Focus Area funded by DOE contract DE-AC02-05CH11231, the Office of Science and Office of Biological and Environmental Research (Lawrence Berkeley National Lab; Operated by the University of California, Berkeley) and National Institutes of Health (NIH) under awards RAI092531A and R01-GM109454, Chan Zuckerberg Biohub and the UC Berkeley-based Innovative Genomics Institute.

## Disclosure declaration

The authors declare that they have no competing interests.

## Notes

#### Summary of Updates

Table 1 revised re suggestions of adding more published CMAGs.

## References

Ackelsberg J, Rakeman J, Hughes S, Petersen J, Mead P, Schriefer M, Kingry L, Hoffmaster A, Gee JE. 2015. Lack of Evidence for Plague or Anthrax on the New York City Subway. Cell Syst 1: 4–5.

Afshinnekoo E, Meydan C, Chowdhury S, Jaroudi D, Boyer C, Bernstein N, Maritz JM, Reeves D, Gandara J, Chhangawala S, et al. 2015. Geospatial Resolution of Human and Bacterial Diversity with City-Scale Metagenomics. Cell Syst 1: 97–97.e3.

Albertsen M, Hugenholtz P, Skarshewski A, Nielsen KL, Tyson GW, Nielsen PH. 2013. Genome sequences of rare, uncultured bacteria obtained by differential coverage binning of multiple metagenomes. Nat Biotechnol 31: 533–538.

Almeida A, Mitchell AL, Boland M, Forster SC, Gloor GB, Tarkowska A, Lawley TD, Finn RD. 2019. A new genomic blueprint of the human gut microbiota. Nature 568: 499–504.

Alneberg J, Bjarnason BS, de Bruijn I, Schirmer M, Quick J, Ijaz UZ, Loman NJ, Andersson AF, Quince C. 2013. CONCOCT: Clustering cONtigs on COverage and ComposiTion. arXiv [q-bioGN]. http://arxiv.org/abs/1312.4038.

Al-Shayeb B, Sachdeva R, Chen LX, Ward F, Munk P. 2019. Clades of huge phage from across Earth’s ecosystems. bioRxiv. https://www.biorxiv.org/content/10.1101/572362v1.abstract.

Altschul SF, Gish W, Miller W, Myers EW, Lipman DJ. 1990. Basic local alignment search tool. J Mol Biol 215: 403–410.

Anantharaman K, Brown CT, Hug LA, Sharon I, Castelle CJ, Probst AJ, Thomas BC, Singh A, Wilkins MJ, Karaoz U, et al. 2016. Thousands of microbial genomes shed light on interconnected biogeochemical processes in an aquifer system. Nature Communications 7. http://dx.doi.org/10.1038/ncomms13219.

Anantharaman K, Hausmann B, Jungbluth SP, Kantor RS, Lavy A, Warren LA, Rappé MS, Pester M, Loy A, Thomas BC, et al. 2018. Expanded diversity of microbial groups that shape the dissimilatory sulfur cycle. ISME J 12: 1715–1728.

Arakawa K. 2016. No evidence for extensive horizontal gene transfer from the draft genome of a tardigrade. Proc Natl Acad Sci U S A 113: E3057.

Arumugam K, Bağcı C, Bessarab I, Beier S, Buchfink B, Górska A, Qiu G, Huson DH, Williams RBH. 2019. Annotated bacterial chromosomes from frame-shift-corrected long-read metagenomic data. Microbiome 7: 61.

Banfield JF, Anantharaman K, Williams KH, Thomas BC. 2017. Complete 4.55-Megabase-Pair Genome of “Candidatus Fluviicola riflensis,” Curated from Short-Read Metagenomic Sequences. Genome Announc 5: e01299–17.

Barry ER, Bell SD. 2006. DNA replication in the archaea. Microbiol Mol Biol Rev 70: 876–887.

Baudry L, Foutel-Rodier T, Thierry A, Koszul R, Marbouty M. 2019. MetaTOR: A Computational Pipeline to Recover High-Quality Metagenomic Bins From Mammalian Gut Proximity-Ligation (meta3C) Libraries. Front Genet 10: 753.

Becraft ED, Woyke T, Jarett J, Ivanova N, Godoy-Vitorino F, Poulton N, Brown JM, Brown J, Lau MCY, Onstott T, et al. 2017. Rokubacteria: Genomic Giants among the Uncultured Bacterial Phyla. Front Microbiol 8: 2264.

Bickhart DM, Watson M, Koren S, Panke-Buisse K, Cersosimo LM, Press MO, Van Tassell CP, Van Kessel JAS, Haley BJ, Kim SW, et al. 2019. Assignment of virus and antimicrobial resistance genes to microbial hosts in a complex microbial community by combined long-read assembly and proximity ligation. Genome Biol 20: 153.

Boothby TC, Tenlen JR, Smith FW, Wang JR, Patanella KA, Nishimura EO, Tintori SC, Li Q, Jones CD, Yandell M, et al. 2015. Evidence for extensive horizontal gene transfer from the draft genome of a tardigrade. Proc Natl Acad Sci U S A 112: 15976–15981.

Bor B, Bedree JK, Shi W, McLean JS, He X. 2019. Saccharibacteria (TM7) in the Human Oral Microbiome. J Dent Res 98: 500–509.

Bowers RM, Kyrpides NC, Stepanauskas R, Harmon-Smith M, Doud D, Reddy TBK, Schulz F, Jarett J, Rivers AR, Eloe-Fadrosh EA, et al. 2017. Minimum information about a single amplified genome (MISAG) and a metagenome-assembled genome (MIMAG) of bacteria and archaea. Nat Biotechnol 35: 725.

Brooks B, Olm MR, Firek BA, Baker R, Thomas BC, Morowitz MJ, Banfield JF. 2017. Strain-resolved analysis of hospital rooms and infants reveals overlap between the human and room microbiome. Nat Commun 8: 1814.

Brown CT, Hug LA, Thomas BC, Sharon I, Castelle CJ, Singh A, Wilkins MJ, Wrighton KC, Williams KH, Banfield JF. 2015. Unusual biology across a group comprising more than 15% of domain Bacteria. Nature 523: 208–211.

Brown CT, Olm MR, Thomas BC, Banfield JF. 2016. Measurement of bacterial replication rates in microbial communities. Nat Biotechnol 34: 1256–1263.

Bushnell B. 2018. BBTools: a suite of fast, multithreaded bioinformatics tools designed for analysis of DNA and RNA sequence data. Joint Genome Institute https://jgidoegov/data-and-tools/bbtools.

Campbell JH, O’Donoghue P, Campbell AG, Schwientek P, Sczyrba A, Woyke T, Söll D, Podar M. 2013. UGA is an additional glycine codon in uncultured SR1 bacteria from the human microbiota. Proc Natl Acad Sci U S A 110: 5540–5545.

Capella-Gutiérrez S, Silla-Martínez JM, Gabaldón T. 2009. trimAl: a tool for automated alignment trimming in large-scale phylogenetic analyses. Bioinformatics 25: 1972–1973.

Castelle CJ, Hug LA, Wrighton KC, Thomas BC, Williams KH, Wu D, Tringe SG, Singer SW, Eisen JA, Banfield JF. 2013. Extraordinary phylogenetic diversity and metabolic versatility in aquifer sediment. Nature Communications 4. http://dx.doi.org/10.1038/ncomms3120.

Chen LX, Zhao YL, McMahon KD, Mori JF, Jessen GL. 2019. Wide distribution of phage that infect freshwater SAR11 bacteria. bioRxiv. https://www.biorxiv.org/content/10.1101/672428v1.abstract.

Chivian D, Brodie EL, Alm EJ, Culley DE, Dehal PS, DeSantis TZ, Gihring TM, Lapidus A, Lin L-H, Lowry SR, et al. 2008. Environmental genomics reveals a single-species ecosystem deep within Earth. Science 322: 275–278.

Daims H, Lebedeva EV, Pjevac P, Han P, Herbold C, Albertsen M, Jehmlich N, Palatinszky M, Vierheilig J, Bulaev A, et al. 2015. Complete nitrification by Nitrospira bacteria. Nature 528: 504.

Dekker J, Rippe K, Dekker M, Kleckner N. 2002. Capturing chromosome conformation. Science 295: 1306–1311.

Delmont TO, Eren AM. 2016. Identifying contamination with advanced visualization and analysis practices: metagenomic approaches for eukaryotic genome assemblies. PeerJ 4: e1839.

Delmont TO, Kiefl E, Kilinc O, Esen OC, Uysal I, Rappé MS, Giovannoni S, Eren AM. 2019. Single-amino acid variants reveal evolutionary processes that shape the biogeography of a global SAR11 subclade. Elife 8. http://dx.doi.org/10.7554/eLife.46497.

Delmont TO, Quince C, Shaiber A, Esen ÖC, Lee ST, Rappé MS, McLellan SL, Lücker S, Eren AM. 2018. Nitrogen-fixing populations of Planctomycetes and Proteobacteria are abundant in surface ocean metagenomes. Nat Microbiol 3: 804–813.

DeMaere MZ, Darling AE. 2019. bin3C: exploiting Hi-C sequencing data to accurately resolve metagenome-assembled genomes. Genome Biology 20. http://dx.doi.org/10.1186/s13059-019-1643-1.

Deorowicz S, Debudaj-Grabysz A, Gudyś A. 2016. FAMSA: Fast and accurate multiple sequence alignment of huge protein families. Sci Rep 6: 33964.

Devoto AE, Santini JM, Olm MR, Anantharaman K, Munk P, Tung J, Archie EA, Turnbaugh PJ, Seed KD, Blekhman R, et al. 2019. Megaphages infect Prevotella and variants are widespread in gut microbiomes. Nat Microbiol 4: 693–700.

Di Rienzi SC, Sharon I, Wrighton KC, Koren O, Hug LA, Thomas BC, Goodrich JK, Bell JT, Spector TD, Banfield JF, et al. 2013. The human gut and groundwater harbor non-photosynthetic bacteria belonging to a new candidate phylum sibling to Cyanobacteria. Elife 2: e01102.

Driscoll CB, Otten TG, Brown NM, Dreher TW. 2017. Towards long-read metagenomics: complete assembly of three novel genomes from bacteria dependent on a diazotrophic cyanobacterium in a freshwater lake co-culture. Standards in Genomic Sciences 12. http://dx.doi.org/10.1186/s40793-017-0224-8.

Eddy SR. 2011. Accelerated Profile HMM Searches. PLoS Comput Biol 7: e1002195.

Eisen JA, Heidelberg JF, White O, Salzberg SL. 2000. Evidence for symmetric chromosomal inversions around the replication origin in bacteria. Genome Biol 1: RESEARCH0011.

Eren AM, Borisy GG, Huse SM, Mark Welch JL. 2014. Oligotyping analysis of the human oral microbiome. Proc Natl Acad Sci U S A 111: E2875–84.

Eren AM, Esen ÖC, Quince C, Vineis JH, Morrison HG, Sogin ML, Delmont TO. 2015. Anvi’o: an advanced analysis and visualization platform for ‘omics data. PeerJ 3: e1319.

Fraser CM, Eisen JA, Nelson KE, Paulsen IT, Salzberg SL. 2002. The value of complete microbial genome sequencing (you get what you pay for). J Bacteriol 184: 6403–6405.

Garcia SL, Stevens SLR, Crary B, Martinez-Garcia M, Stepanauskas R, Woyke T, Tringe SG, Andersson SGE, Bertilsson S, Malmstrom RR, et al. 2018. Contrasting patterns of genome-level diversity across distinct co-occurring bacterial populations. ISME J 12: 742–755.

Garg SG, Kapust N, Lin W, Tria FDK, Nelson-Sathi S, Gould SB, Fan L, Zhu R, Zhang C, Martin WF. 2019. Anomalous phylogenetic behavior of ribosomal proteins in metagenome assembled genomes. bioRxiv 731091. https://www.biorxiv.org/content/10.1101/731091v1.abstract (Accessed August 15, 2019).

Giovannoni SJ. 2017. SAR11 Bacteria: The Most Abundant Plankton in the Oceans. Ann Rev Mar Sci 9: 231–255.

Greenwald WW, Klitgord N, Seguritan V, Yooseph S, Venter JC, Garner C, Nelson KE, Li W. 2017. Utilization of defined microbial communities enables effective evaluation of meta-genomic assemblies. BMC Genomics 18: 296.

Grigoriev A. 1998. Analyzing genomes with cumulative skew diagrams. Nucleic Acids Res 26: 2286–2290.

He X, McLean JS, Edlund A, Yooseph S, Hall AP, Liu S-Y, Dorrestein PC, Esquenazi E, Hunter RC, Cheng G, et al. 2015. Cultivation of a human-associated TM7 phylotype reveals a reduced genome and epibiotic parasitic lifestyle. Proc Natl Acad Sci U S A 112: 244–249.

Hongoh Y, Sharma VK, Prakash T, Noda S, Taylor TD, Kudo T, Sakaki Y, Toyoda A, Hattori M, Ohkuma M. 2008. Complete genome of the uncultured Termite Group 1 bacteria in a single host protist cell. Proc Natl Acad Sci U S A 105: 5555–5560.

Hug LA, Baker BJ, Anantharaman K, Brown CT, Probst AJ, Castelle CJ, Butterfield CN, Hernsdorf AW, Amano Y, Ise K, et al. 2016. A new view of the tree of life. Nat Microbiol 1: 16048.

Hyatt D, Chen G-L, Locascio PF, Land ML, Larimer FW, Hauser LJ. 2010. Prodigal: prokaryotic gene recognition and translation initiation site identification. BMC Bioinformatics 11: 119.

Iverson V, Morris RM, Frazar CD, Berthiaume CT, Morales RL, Armbrust EV. 2012. Untangling genomes from metagenomes: revealing an uncultured class of marine Euryarchaeota. Science 335: 587–590.

Kalisky T, Quake SR. 2011. Single-cell genomics. Nat Methods 8: 311–314.

Kang DD, Froula J, Egan R, Wang Z. 2015. MetaBAT, an efficient tool for accurately reconstructing single genomes from complex microbial communities. PeerJ 3: e1165.

Kantor RS, Huddy RJ, Iyer R, Thomas BC, Brown CT, Anantharaman K, Tringe S, Hettich RL, Harrison STL, Banfield JF. 2017. Genome-Resolved Meta-Omics Ties Microbial Dynamics to Process Performance in Biotechnology for Thiocyanate Degradation. Environ Sci Technol 51: 2944–2953.

Kantor RS, Wrighton KC, Handley KM, Sharon I, Hug LA, Castelle CJ, Thomas BC, Banfield JF. 2013. Small genomes and sparse metabolisms of sediment-associated bacteria from four candidate phyla. MBio 4: e00708–13.

Kashtan N, Roggensack SE, Rodrigue S, Thompson JW, Biller SJ, Coe A, Ding H, Marttinen P, Malmstrom RR, Stocker R, et al. 2014. Single-cell genomics reveals hundreds of coexisting subpopulations in wild Prochlorococcus. Science 344: 416–420.

Kearse M, Moir R, Wilson A, Stones-Havas S, Cheung M, Sturrock S, Buxton S, Cooper A, Markowitz S, Duran C, et al. 2012. Geneious Basic: an integrated and extendable desktop software platform for the organization and analysis of sequence data. Bioinformatics 28: 1647–1649.

Korem T, Zeevi D, Suez J, Weinberger A, Avnit-Sagi T, Pompan-Lotan M, Matot E, Jona G, Harmelin A, Cohen N, et al. 2015. Growth dynamics of gut microbiota in health and disease inferred from single metagenomic samples. Science 349: 1101–1106.

Köster J, Rahmann S. 2012. Snakemake--a scalable bioinformatics workflow engine. Bioinformatics 28: 2520– 2522.

Kowarsky M, Camunas-Soler J, Kertesz M, De Vlaminck I, Koh W, Pan W, Martin L, Neff NF, Okamoto J, Wong RJ, et al. 2017. Numerous uncharacterized and highly divergent microbes which colonize humans are revealed by circulating cell-free DNA. Proc Natl Acad Sci U S A. http://dx.doi.org/10.1073/pnas.1707009114.

Labonté JM, Swan BK, Poulos B, Luo H, Koren S, Hallam SJ, Sullivan MB, Woyke T, Eric Wommack K, Stepanauskas R. 2015. Single-cell genomics-based analysis of virus–host interactions in marine surface bacterioplankton. ISME J 9: 2386–2399.

Laczny CC, Sternal T, Plugaru V, Gawron P, Atashpendar A, Margossian HH, Coronado S, van der Maaten L, Vlassis N, Wilmes P. 2015. VizBin - an application for reference-independent visualization and human-augmented binning of metagenomic data. Microbiome 3: 1.

Langmead B, Salzberg SL. 2012. Fast gapped-read alignment with Bowtie 2. Nat Methods 9: 357–359.

Laver T, Harrison J, O’Neill PA, Moore K, Farbos A, Paszkiewicz K, Studholme DJ. 2015. Assessing the performance of the Oxford Nanopore Technologies MinION. Biomol Detect Quantif 3: 1–8.

Le TBK, Imakaev MV, Mirny LA, Laub MT. 2013. High-resolution mapping of the spatial organization of a bacterial chromosome. Science 342: 731–734.

Lieberman-Aiden E, van Berkum NL, Williams L, Imakaev M, Ragoczy T, Telling A, Amit I, Lajoie BR, Sabo PJ, Dorschner MO, et al. 2009. Comprehensive mapping of long-range interactions reveals folding principles of the human genome. Science 326: 289–293.

Li H, Handsaker B, Wysoker A, Fennell T, Ruan J, Homer N, Marth G, Abecasis G, Durbin R, 1000 Genome Project Data Processing Subgroup. 2009. The Sequence Alignment/Map format and SAMtools. Bioinformatics 25: 2078–2079.

Liu M, Darling A. 2015. Metagenomic Chromosome Conformation Capture (3C): techniques, applications, and challenges. F1000Res 4: 1377.

Lobry JR. 1996. Origin of replication of Mycoplasma genitalium. Science 272: 745–746.

Lo I, Denef VJ, Verberkmoes NC, Shah MB, Goltsman D, DiBartolo G, Tyson GW, Allen EE, Ram RJ, Detter JC, et al. 2007. Strain-resolved community proteomics reveals recombining genomes of acidophilic bacteria. Nature 446: 537–541.

Lowe TM, Chan PP. 2016. tRNAscan-SE On-line: integrating search and context for analysis of transfer RNA genes. Nucleic Acids Res 44: W54–7.

Luef B, Frischkorn KR, Wrighton KC, Holman H-YN, Birarda G, Thomas BC, Singh A, Williams KH, Siegerist CE, Tringe SG, et al. 2015. Diverse uncultivated ultra-small bacterial cells in groundwater. Nat Commun 6: 6372.

Marbouty M, Cournac A, Flot J-F, Marie-Nelly H, Mozziconacci J, Koszul R. 2014. Metagenomic chromosome conformation capture (meta3C) unveils the diversity of chromosome organization in microorganisms. Elife 3: e03318.

Marbouty M, Koszul R. 2015. Metagenome Analysis Exploiting High-Throughput Chromosome Conformation Capture (3C) Data. Trends in Genetics 31: 673–682. http://dx.doi.org/10.1016/j.tig.2015.10.003.

Mark Welch JL, Utter DR, Rossetti BJ, Mark Welch DB, Eren AM, Borisy GG. 2014. Dynamics of tongue microbial communities with single-nucleotide resolution using oligotyping. Front Microbiol 5: 568.

Mosier AC, Miller CS, Frischkorn KR, Ohm RA, Li Z, LaButti K, Lapidus A, Lipzen A, Chen C, Johnson J, et al. 2016. Fungi Contribute Critical but Spatially Varying Roles in Nitrogen and Carbon Cycling in Acid Mine Drainage. Frontiers in Microbiology 7. http://dx.doi.org/10.3389/fmicb.2016.00238.

Moustafa A, Xie C, Kirkness E, Biggs W, Wong E, Turpaz Y, Bloom K, Delwart E, Nelson KE, Venter JC, et al. 2017. The blood DNA virome in 8,000 humans. PLoS Pathog 13: e1006292.

Nadalin F, Vezzi F, Policriti A. 2012. GapFiller: a de novo assembly approach to fill the gap within paired reads. BMC Bioinformatics 13 Suppl 14: S8.

Nakamura Y. 2002. Complete Genome Structure of the Thermophilic Cyanobacterium Thermosynechococcus elongatus BP-1. DNA Research 9: 123–130. http://dx.doi.org/10.1093/dnares/9.4.123.

Nayfach S, Shi ZJ, Seshadri R, Pollard KS, Kyrpides N. 2019. Novel insights from uncultivated genomes of the global human gut microbiome. Nature. http://dx.doi.org/10.1038/s41586-019-1058-x.

Nelson WC, Stegen JC. 2015. The reduced genomes of Parcubacteria (OD1) contain signatures of a symbiotic lifestyle. Front Microbiol 6: 713.

Newton RJ, Jones SE, Helmus MR, McMahon KD. 2007. Phylogenetic ecology of the freshwater Actinobacteria acI lineage. Appl Environ Microbiol 73: 7169–7176.

Nguyen L-T, Schmidt HA, von Haeseler A, Minh BQ. 2015. IQ-TREE: a fast and effective stochastic algorithm for estimating maximum-likelihood phylogenies. Mol Biol Evol 32: 268–274.

Nicholls SM, Quick JC, Tang S, Loman NJ. 2019. Ultra-deep, long-read nanopore sequencing of mock microbial community standards. Gigascience 8. http://dx.doi.org/10.1093/gigascience/giz043.

Nurk S, Meleshko D, Korobeynikov A, Pevzner PA. 2017. metaSPAdes: a new versatile metagenomic assembler. Genome Res 27: 824–834.

Olm MR, Brown CT, Brooks B, Firek B, Baker R, Burstein D, Soenjoyo K, Thomas BC, Morowitz M, Banfield JF. 2017. Identical bacterial populations colonize premature infant gut, skin, and oral microbiomes and exhibit different in situ growth rates. Genome Res 27: 601–612.

Olm MR, West PT, Brooks B, Firek BA, Baker R, Morowitz MJ, Banfield JF. 2019. Genome-resolved metagenomics of eukaryotic populations during early colonization of premature infants and in hospital rooms. Microbiome 7: 26.

Olson ND, Treangen TJ, Hill CM, Cepeda-Espinoza V, Ghurye J, Koren S, Pop M. 2017. Metagenomic assembly through the lens of validation: recent advances in assessing and improving the quality of genomes assembled from metagenomes. Brief Bioinform. http://dx.doi.org/10.1093/bib/bbx098.

Parks DH, Chuvochina M, Waite DW, Rinke C, Skarshewski A, Chaumeil P-A, Hugenholtz P. 2018. A standardized bacterial taxonomy based on genome phylogeny substantially revises the tree of life. Nat Biotechnol 36: 996–1004.

Parks DH, Imelfort M, Skennerton CT, Hugenholtz P, Tyson GW. 2015. CheckM: assessing the quality of microbial genomes recovered from isolates, single cells, and metagenomes. Genome Res 25: 1043–1055.

Parks DH, Rinke C, Chuvochina M, Chaumeil P-A, Woodcroft BJ, Evans PN, Hugenholtz P, Tyson GW. 2017. Recovery of nearly 8,000 metagenome-assembled genomes substantially expands the tree of life. Nat Microbiol 2: 1533–1542.

Pasolli E, Asnicar F, Manara S, Zolfo M, Karcher N, Armanini F, Beghini F, Manghi P, Tett A, Ghensi P, et al. 2019. >Extensive Unexplored Human Microbiome Diversity Revealed by Over 150,000 Genomes from Metagenomes Spanning Age, Geography, and Lifestyle. Cell 176: 649–662.e20.

Pelletier E, Kreimeyer A, Bocs S, Rouy Z, Gyapay G, Chouari R, Rivière D, Ganesan A, Daegelen P, Sghir A, et al. 2008. “Candidatus Cloacamonas Acidaminovorans”: Genome Sequence Reconstruction Provides a First Glimpse of a New Bacterial Division. J Bacteriol 190: 2572–2579.

Peng Y, Leung HCM, Yiu SM, Chin FYL. 2012. IDBA-UD: a de novo assembler for single-cell and metagenomic sequencing data with highly uneven depth. Bioinformatics 28: 1420–1428.

Probst AJ, Banfield JF. 2018. Homologous Recombination and Transposon Propagation Shape the Population Structure of an Organism from the Deep Subsurface with Minimal Metabolism. Genome Biol Evol 10: 1115–1119.

Probst AJ, Ladd B, Jarett JK, Geller-McGrath DE, Sieber CMK, Emerson JB, Anantharaman K, Thomas BC, Malmstrom RR, Stieglmeier M, et al. 2018. Differential depth distribution of microbial function and putative symbionts through sediment-hosted aquifers in the deep terrestrial subsurface. Nat Microbiol 3: 328–336.

Quandt CA, Alisha Quandt C, Kohler A, Hesse CN, Sharpton TJ, Martin F, Spatafora JW. 2015. Metagenome sequence of Elaphomyces granulatus from sporocarp tissue reveals Ascomycota ectomycorrhizal fingerprints of genome expansion and aProteobacteria-rich microbiome. Environmental Microbiology 17: 2952–2968. http://dx.doi.org/10.1111/1462-2920.12840.

Quince C, Delmont TO, Raguideau S, Alneberg J, Darling AE, Collins G, Eren AM. 2017. DESMAN: a new tool for de novo extraction of strains from metagenomes. Genome Biol 18: 181.

Rang FJ, Kloosterman WP, de Ridder J. 2018. From squiggle to basepair: computational approaches for improving nanopore sequencing read accuracy. Genome Biol 19: 90.

Rao SSP, Huntley MH, Durand NC, Stamenova EK, Bochkov ID, Robinson JT, Sanborn AL, Machol I, Omer AD, Lander ES, et al. 2014. A 3D map of the human genome at kilobase resolution reveals principles of chromatin looping. Cell 159: 1665–1680.

Raveh-Sadka T, Thomas BC, Singh A, Firek B, Brooks B, Castelle CJ, Sharon I, Baker R, Good M, Morowitz MJ, et al. 2015. Gut bacteria are rarely shared by co-hospitalized premature infants, regardless of necrotizing enterocolitis development. Elife 4. http://dx.doi.org/10.7554/eLife.05477.

Reveillaud J, Bordenstein SR, Cruaud C, Shaiber A, Esen ÖC, Weill M, Makoundou P, Lolans K, Watson AR, Rakotoarivony I, et al. 2019. The Wolbachia mobilome in Culex pipiens includes a putative plasmid. Nat Commun 10: 1051.

Rinke C, Schwientek P, Sczyrba A, Ivanova NN, Anderson IJ, Cheng J-F, Darling A, Malfatti S, Swan BK, Gies EA, et al. 2013. Insights into the phylogeny and coding potential of microbial dark matter. Nature 499: 431–437.

Robinson JT, Thorvaldsdóttir H, Winckler W, Guttman M, Lander ES, Getz G, Mesirov JP. 2011. Integrative genomics viewer. Nat Biotechnol 29: 24.

Rocha EP, Danchin A, Viari A. 1999. Universal replication biases in bacteria. Mol Microbiol 32: 11–16.

Rojas-Carulla M, Ley RE, Schölkopf B, Youngblut ND. 2019. DeepMAsED: Evaluating the quality of metagenomic assemblies. http://dx.doi.org/10.1101/763813.

Roux S, Enault F, Hurwitz BL, Sullivan MB. 2015. VirSorter: mining viral signal from microbial genomic data. PeerJ 3: e985.

Sangwan N, Xia F, Gilbert JA. 2016. Recovering complete and draft population genomes from metagenome datasets. Microbiome 4: 8.

Schirmer M, D’Amore R, Ijaz UZ, Hall N, Quince C. 2016. Illumina error profiles: resolving fine-scale variation in metagenomic sequencing data. BMC Bioinformatics 17: 125.

Shaiber A, Eren AM. 2019. Composite Metagenome-Assembled Genomes Reduce the Quality of Public Genome Repositories. MBio 10. http://dx.doi.org/10.1128/mBio.00725-19.

Sharon I, Banfield JF. 2013. Genomes from Metagenomics. Science 342: 1057–1058.

Sharon I, Morowitz MJ, Thomas BC, Costello EK, Relman DA, Banfield JF. 2013. Time series community genomics analysis reveals rapid shifts in bacterial species, strains, and phage during infant gut colonization. Genome Research 23: 111–120. http://dx.doi.org/10.1101/gr.142315.112.

Sieber CMK, Probst AJ, Sharrar A, Thomas BC, Hess M, Tringe SG, Banfield JF. 2018. Recovery of genomes from metagenomes via a dereplication, aggregation and scoring strategy. Nat Microbiol 3: 836–843.

Simmons SL, Dibartolo G, Denef VJ, Goltsman DSA, Thelen MP, Banfield JF. 2008. Population genomic analysis of strain variation in Leptospirillum group II bacteria involved in acid mine drainage formation. PLoS Biol 6: e177.

Stalder T, Press MO, Sullivan S, Liachko I, Top EM. 2019. Linking the resistome and plasmidome to the microbiome. ISME J 13: 2437–2446.

Stepanauskas R. 2012. Single cell genomics: an individual look at microbes. Curr Opin Microbiol 15: 613–620.

Swan BK, Martinez-Garcia M, Preston CM, Sczyrba A, Woyke T, Lamy D, Reinthaler T, Poulton NJ, Masland EDP, Gomez ML, et al. 2011. Potential for chemolithoautotrophy among ubiquitous bacteria lineages in the dark ocean. Science 333: 1296–1300.

Tatusov RL, Galperin MY, Natale DA, Koonin EV. 2000. The COG database: a tool for genome-scale analysis of protein functions and evolution. Nucleic Acids Res 28: 33–36.

Tully BJ, Graham ED, Heidelberg JF. 2018. The reconstruction of 2,631 draft metagenome-assembled genomes from the global oceans. Sci Data 5: 170203.

Turnbaugh PJ, Ley RE, Hamady M, Fraser-Liggett CM, Knight R, Gordon JI. 2007. The human microbiome project. Nature 449: 804–810.

Tyson GW, Chapman J, Hugenholtz P, Allen EE, Ram RJ, Richardson PM, Solovyev VV, Rubin EM, Rokhsar DS, Banfield JF. 2004. Community structure and metabolism through reconstruction of microbial genomes from the environment. Nature 428: 37–43.

Uritskiy GV, DiRuggiero J, Taylor J. 2018. MetaWRAP—a flexible pipeline for genome-resolved metagenomic data analysis. Microbiome 6: 158.

van Kessel MAHJ, Speth DR, Albertsen M, Nielsen PH, Op den Camp HJM, Kartal B, Jetten MSM, Lücker S. 2015. Complete nitrification by a single microorganism. Nature 528: 555–559.

Vineis JH, Ringus DL, Morrison HG, Delmont TO, Dalal S, Raffals LH, Antonopoulos DA, Rubin DT, Eren AM, Chang EB, et al. 2016. Patient-Specific Bacteroides Genome Variants in Pouchitis. MBio 7. http://dx.doi.org/10.1128/mBio.01713-16.

West PT, Probst AJ, Grigoriev IV, Thomas BC, Banfield JF. 2018. Genome-reconstruction for eukaryotes from complex natural microbial communities. Genome Res 28: 569–580.

Whelan S, Goldman N. 2001. A general empirical model of protein evolution derived from multiple protein families using a maximum-likelihood approach. Mol Biol Evol 18: 691–699.

White RA 3rd, Bottos EM, Roy Chowdhury T, Zucker JD, Brislawn CJ, Nicora CD, Fansler SJ, Glaesemann KR, Glass K, Jansson JK. 2016. Moleculo Long-Read Sequencing Facilitates Assembly and Genomic Binning from Complex Soil Metagenomes. mSystems 1. http://dx.doi.org/10.1128/mSystems.00045-16.

Wick RR, Judd LM, Gorrie CL, Holt KE. 2017. Unicycler: Resolving bacterial genome assemblies from short and long sequencing reads. PLoS Comput Biol 13: e1005595.

Woyke T, Doud DFR, Schulz F. 2017. The trajectory of microbial single-cell sequencing. Nat Methods 14: 1045–1054.

Woyke T, Tighe D, Mavromatis K, Clum A, Copeland A, Schackwitz W, Lapidus A, Wu D, McCutcheon JP, McDonald BR, et al. 2010. One Bacterial Cell, One Complete Genome. PLoS ONE 5: e10314. http://dx.doi.org/10.1371/journal.pone.0010314.

Wrighton KC, Thomas BC, Sharon I, Miller CS, Castelle CJ, VerBerkmoes NC, Wilkins MJ, Hettich RL, Lipton MS, Williams KH, et al. 2012. Fermentation, hydrogen, and sulfur metabolism in multiple uncultivated bacterial phyla. Science 337: 1661–1665.

Wu D, Hugenholtz P, Mavromatis K, Pukall R, Dalin E, Ivanova NN, Kunin V, Goodwin L, Wu M, Tindall BJ, et al. 2009. A phylogeny-driven genomic encyclopaedia of Bacteria and Archaea. Nature 462: 1056–1060.

Wu Y-W, Tang Y-H, Tringe SG, Simmons BA, Singer SW. 2014. MaxBin: an automated binning method to recover individual genomes from metagenomes using an expectation-maximization algorithm. Microbiome 2: 26.

