## Supplementary Figures for "Accurate and Complete Genomes from Metagenomes"

**Supplementary Information**
for

**Accurate and Complete Genomes from Metagenomes**

Lin-Xing Chen[1], Karthik Anantharaman[1,7], Alon Shaiber[2,3], A. Murat Eren[3,4,]*, and Jillian F. Banfield[1,5,6]*

[1] Department of Earth and Planetary Sciences, Berkeley, CA, USA.

[2] Graduate Program in Biophysical Sciences, University of Chicago, Chicago, IL 60637, USA.

[3] Department of Medicine, University of Chicago, Chicago 60637 IL, USA.

[4] Bay Paul Center, Marine Biological Laboratory, Woods Hole 02543 MA, USA.

[5] Department of Environmental Science, Policy, and Management, Berkeley, CA, USA.

[6] Earth and Environmental Sciences, Lawrence Berkeley National Laboratory, Berkeley, CA, USA.

[7] Present address: Department of Bacteriology, University of Wisconsin, Madison, WI, USA.

*Corresponding authors:

Address: McCone Hall, Berkeley, CA 94720

Address: 900 E. 57th St., Chicago, IL 60637 USA

**Running title: Curated and complete metagenome-assembled genomes**

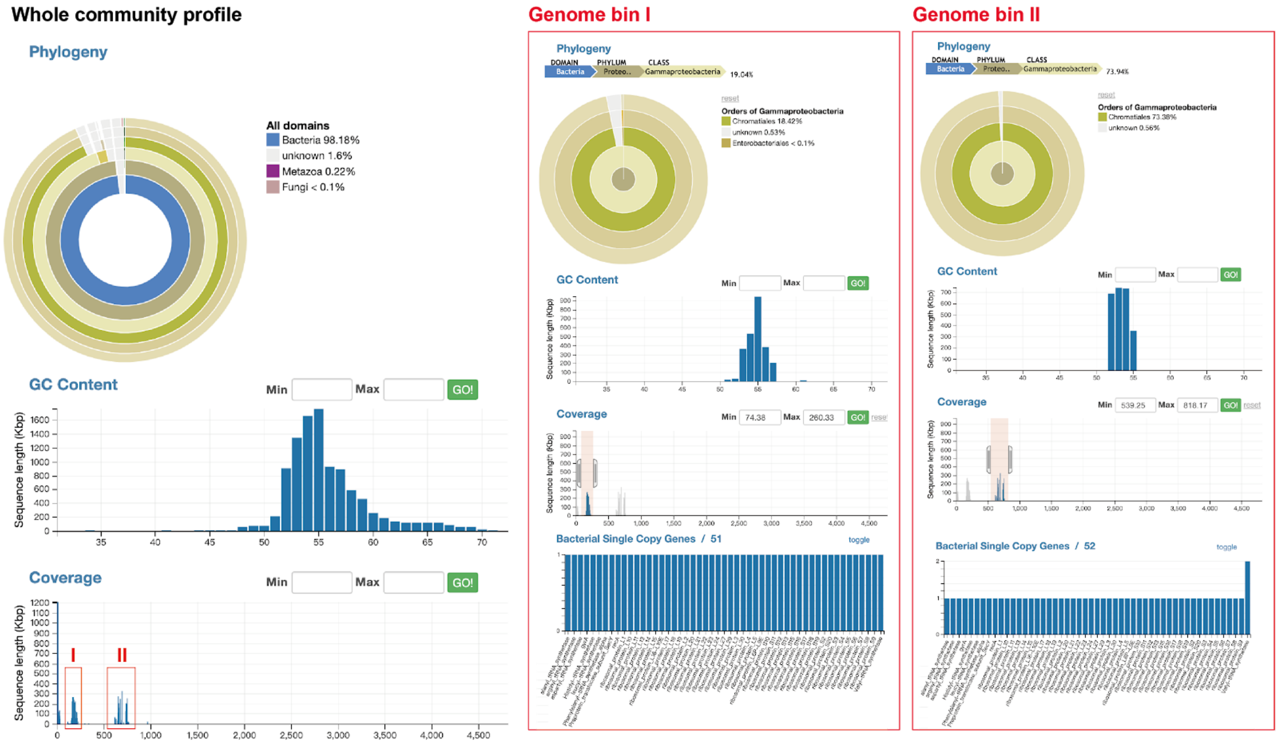


**Figure S1**. Example of bin establishment from a simple microbial community. The two most abundant genomes share similar GC content but the coverage profiles differ so the scaffolds can be easily separated based on coverage. The phylogenetic profile, GC content of the two sets of contigs with the selected coverage ranges and inventory of expected single copy genes are shown on the right.

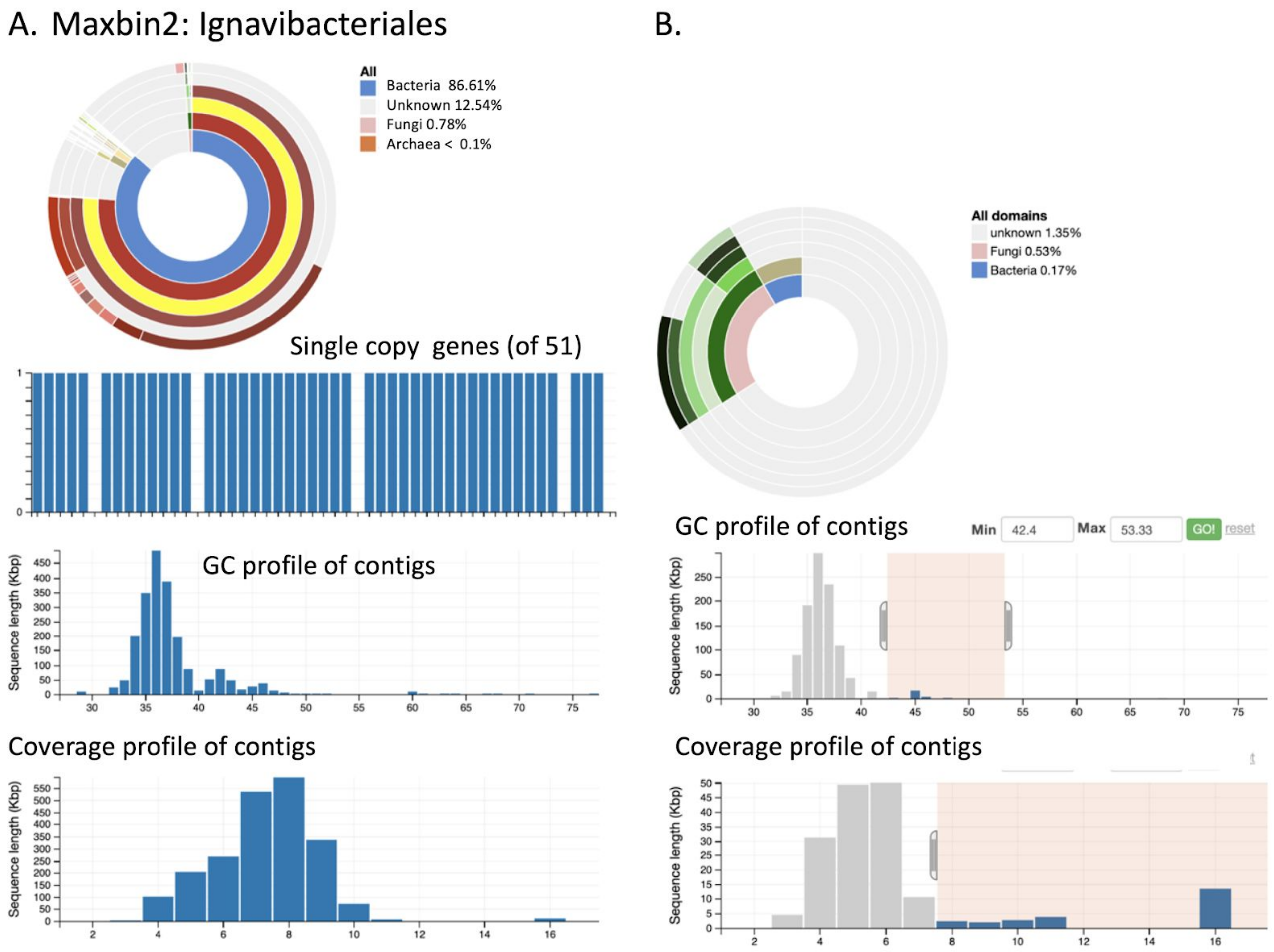


**Figure S2**. Overview of a draft genome generated by autobinning. The DASTool algorithm selected a bin made using Maxbin2. However, like many automatically generated bins, the draft genome contains contaminating sequences that are not evident based on the single copy gene inventory. **A.** Phylogenetic profile (wheel profiling overall contig set at various taxonomic levels, from Domain (inner ring) to species (outermost ring)). Note the dominance of the bin by sequences profiled as Ignavibacteriales but the presence in the bin of sequences from other phyla and domains and without a domain profile (Unknown), peaks with anomalously high GC and some sequence with higher than normal coverage. A 56 kbp phage sequence was removed from the Unknown domain **B.** One step in bin curation: selection of a subset of scaffolds with anomalous coverage and GC content indicates the presence in the bin of eukaryotic and other contaminants.

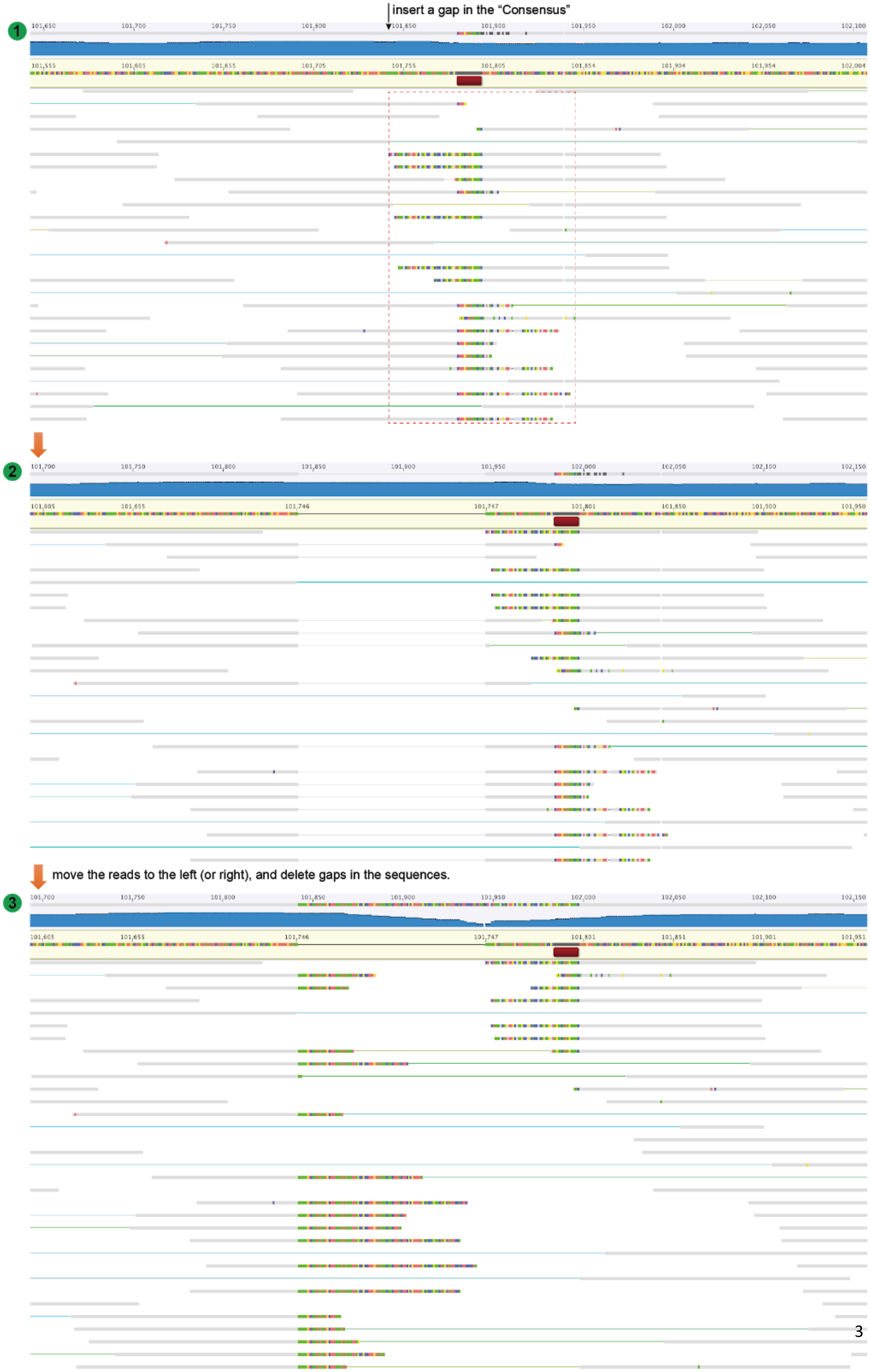

insert a gap in the "Consensus"
1
2
move the reads to the left (or right), and delete gaps in the sequences.
3

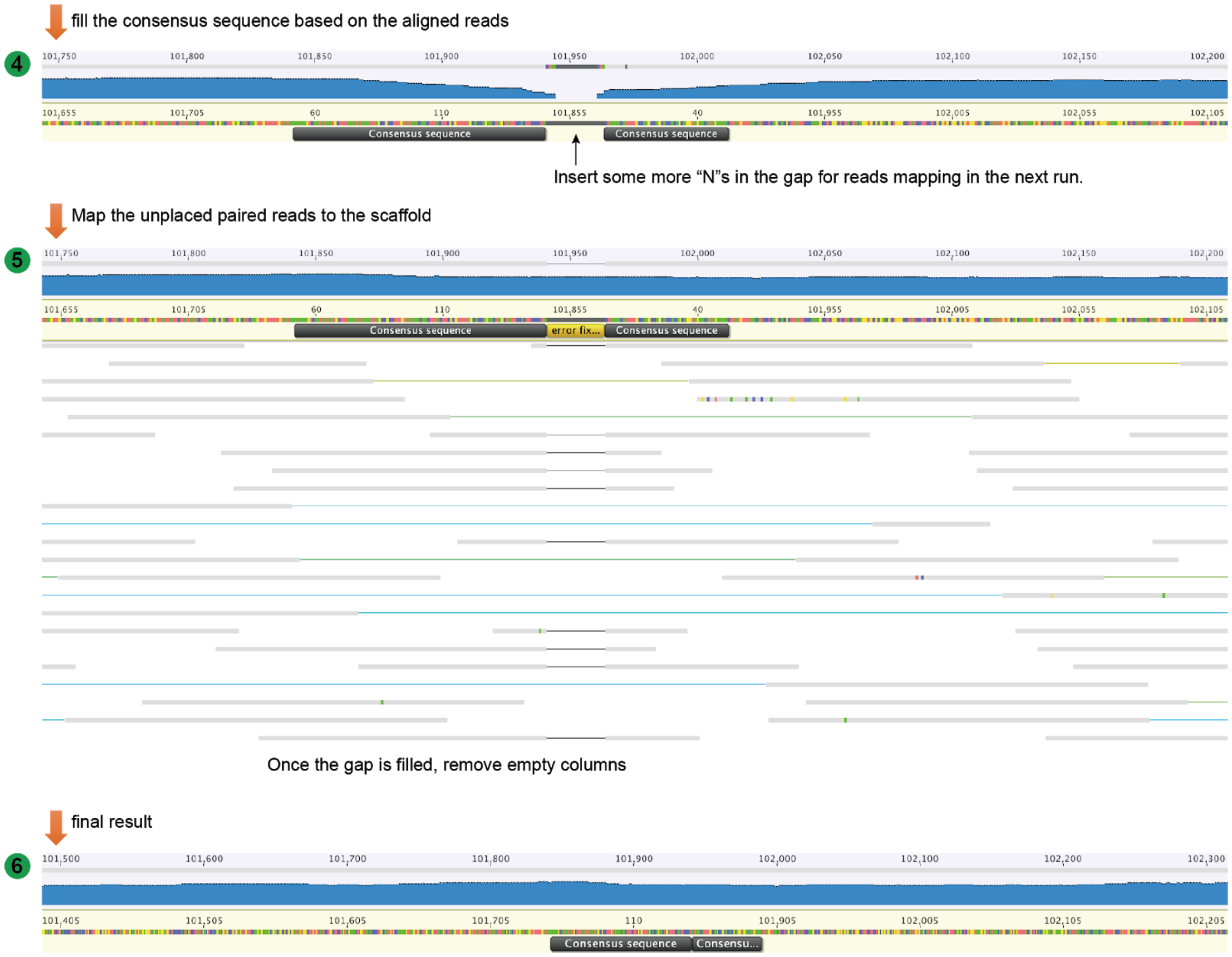


**Note: sometimes you may have to do several cycles of steps 2-4, before the gap is filled.**

**Figure S3.** Diagrams showing the procedure used to fix a local assembly error using uplaced paired-end reads mapped to the problem region after opening a gap.

Ns were wrongly inserted during scaffolding. The sequences flanking the Ns are the same, so the Ns can be deleted and duplicated sequence removed.

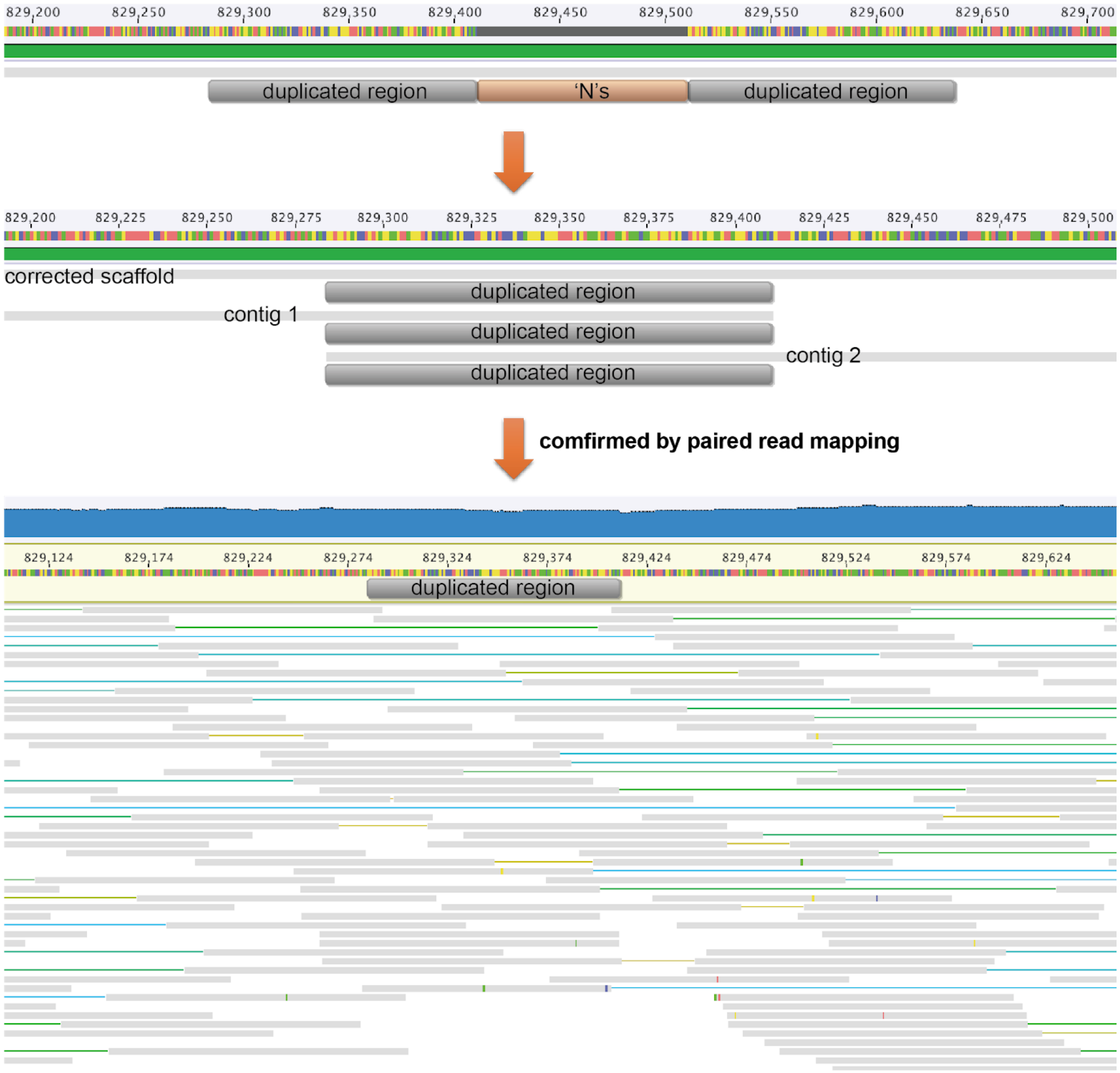


**Figure S4.** The diagram shows a local error where Ns were wrongly inserted during the scaffolding step.

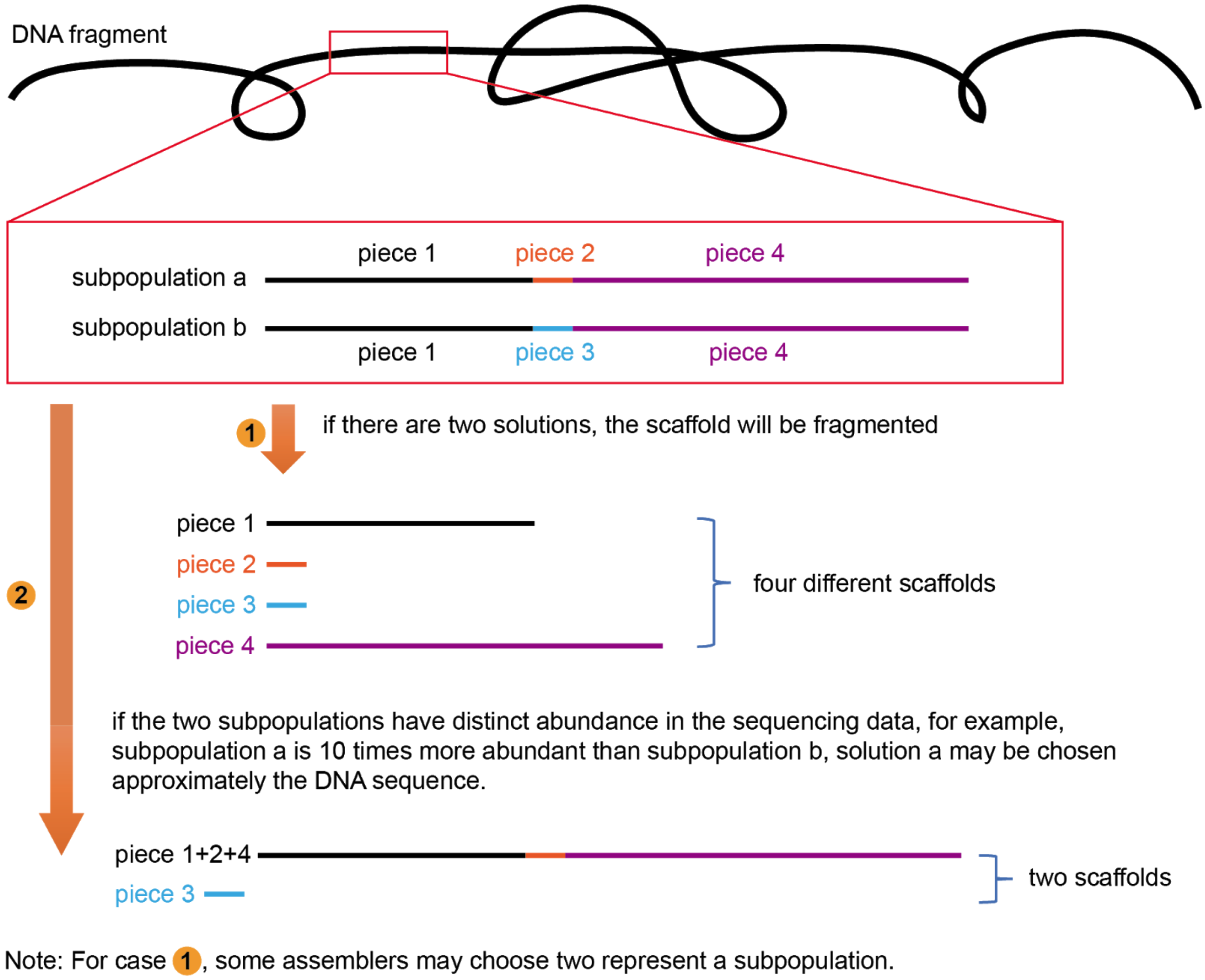


**Figure S5A.** A possible solution when a scaffold breaks during assembly due to subpopulation variation.

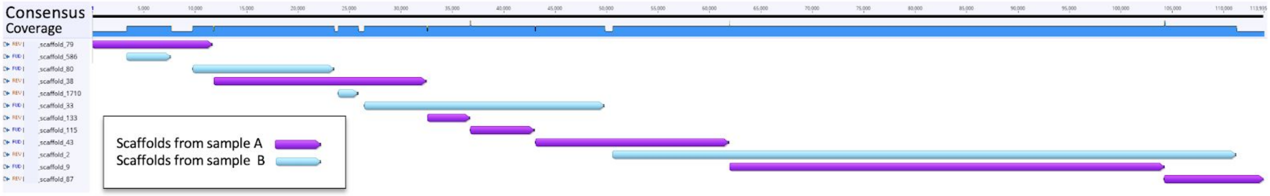


Other examples:

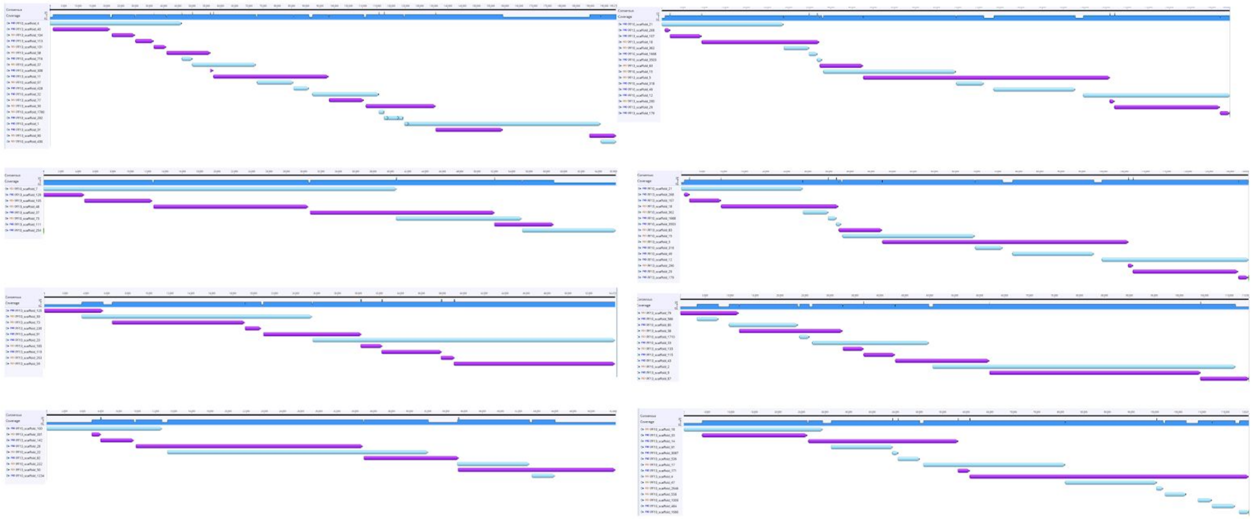

**Figure S5B**. In some cases, scaffold joins may be informed by assembly of the same (or a very closely related) genome in another related sample (for example, from another time/depth/treatment). As shown below, both assembly paths are informed by the assembly of the same genome in another sample. Scaffold joins made based on data from another samples should be treated as hypotheses, and the entire genome verified by read mapping from a single sample. Different fragmentation patterns can occur due to differences in population structure between the samples or differences in coverage.

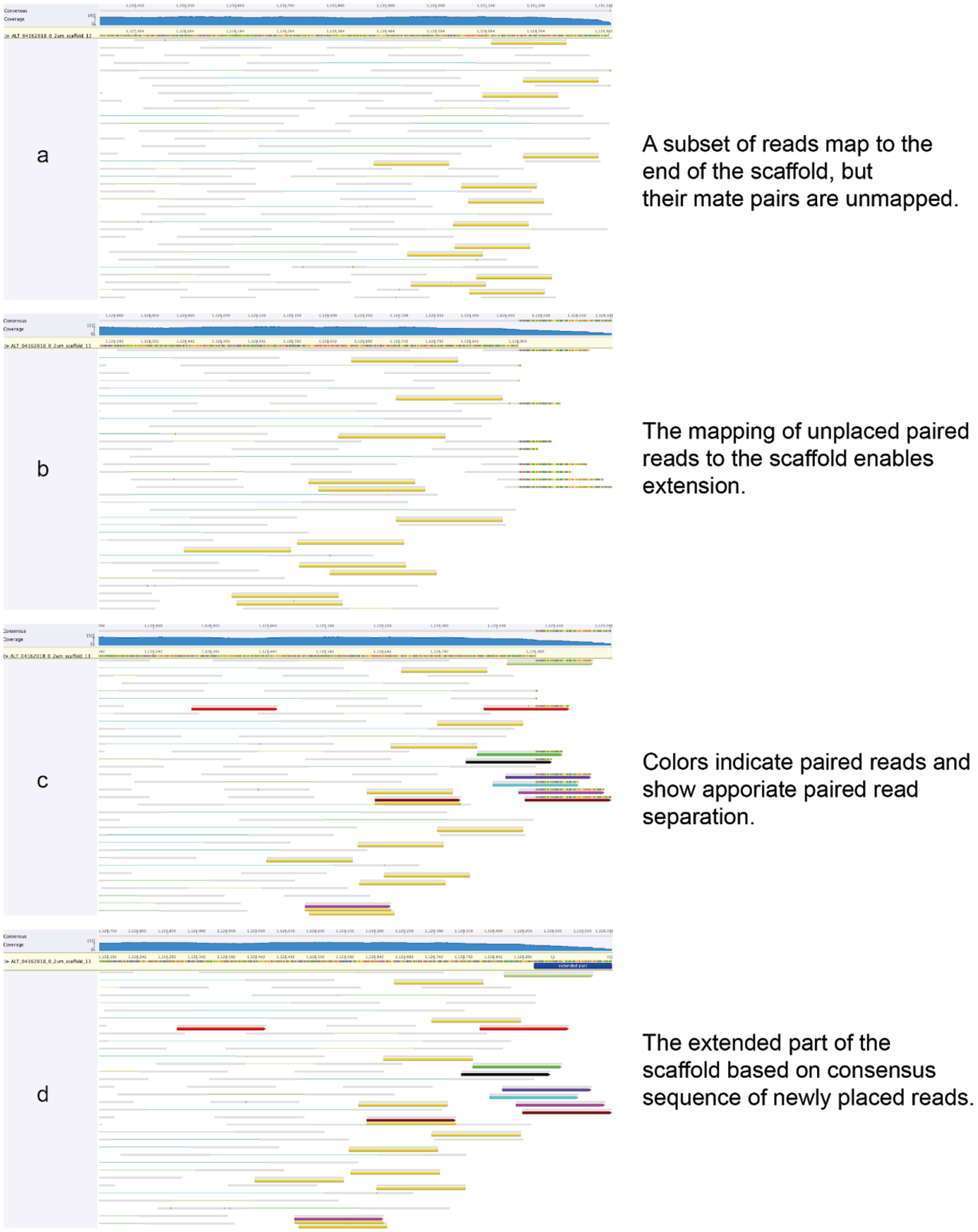


**Figure S6.** The diagram shows the extension of the scaffold end by mapping paired-end reads to it. Extension could be performed for several cycles. Sometimes searching the extended sequence against the whole metagenome can bring in a missing fragment to substantially extend the scaffold. Joins are checked by read mapping.

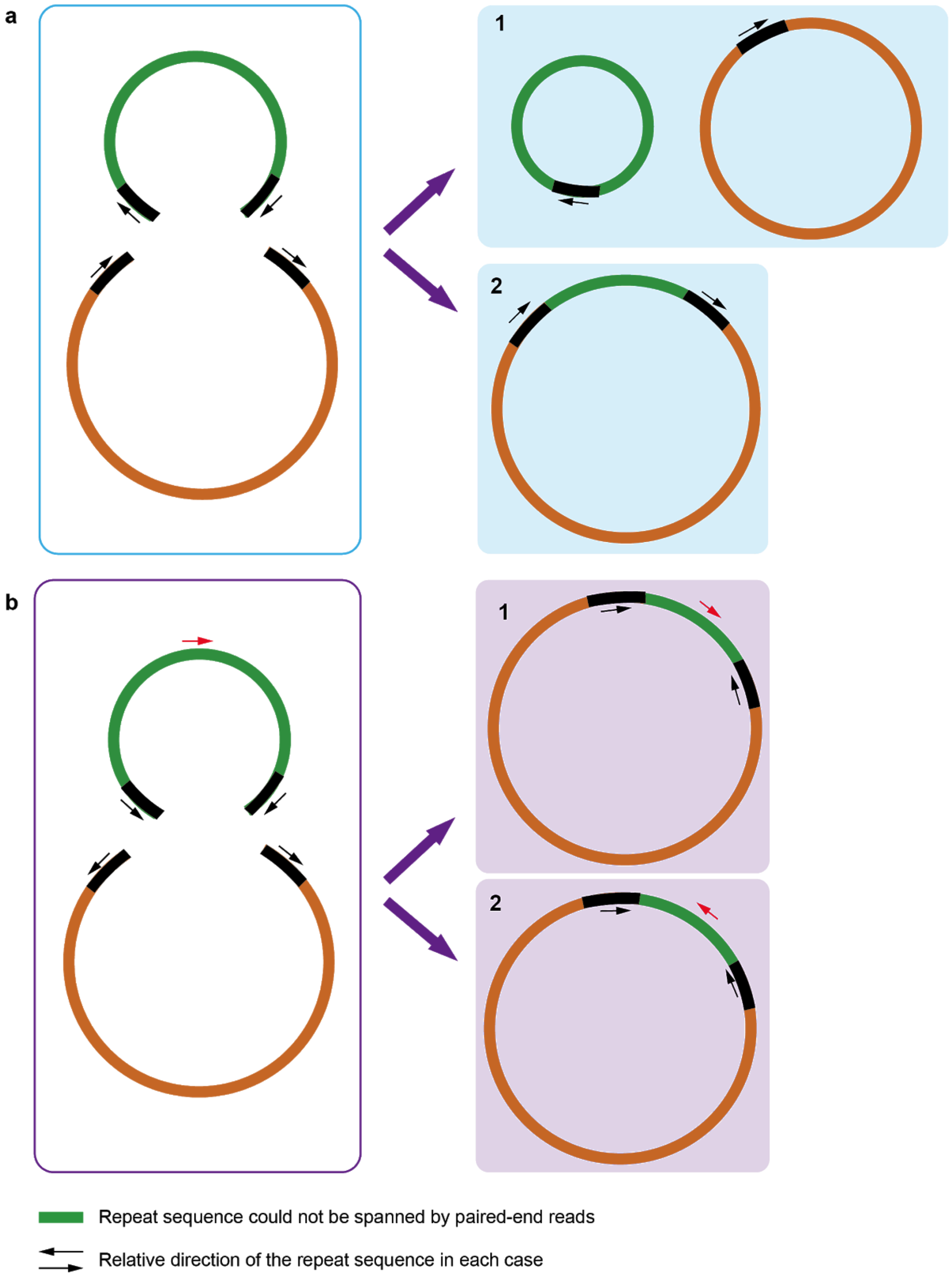


**Figure S7.** Diagrams showing two possible solutions for two cases where a genome is in two pieces and there is no direct solution due to large repeat sequences that could not be spanned by paired-end reads. A. and B. differ in the orientations of the repeated sequences (see arrows). In case A, resolution into a single circular chromosome may be most likely. In case B., the options may be distinguished using GC skew information.

**Figure S8**. ***Case one, a CPR genome curated to completion***. In the step-by-step procedures shown below, we first obtained the mapped reads file for ALT_04162018_0_2um_scaffold_13 (hereafter referred to as "ALT_scaffold_13"). The graph of the coverage of paired reads mapped to ALT_scaffold_13 is shown below.

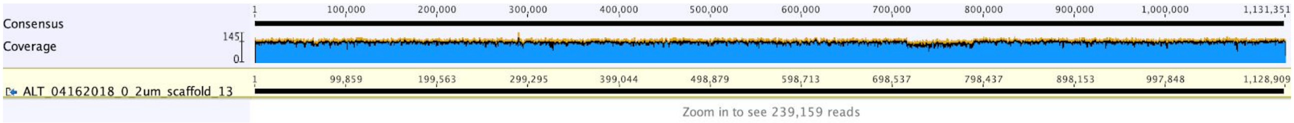


The ends of a scaffold could be extended as shown in Figure S6. After the extension of the scaffold, we searched the extended parts of against the whole metagenome scaffold set to retrieve fragments missed from the genome using BLASTn. Short pieces were identified that link the two ends of ALT_scaffold_13. However, the solutions were not unique due to the presence of SNVs. MetaSPades chose one path whereas IDBA_UD broke the assembly at this position.

***Confirmation of circularization***: The two ends of the scaffold shared the same sequence. We mapped reads to the scaffold (after trimming duplicated sequence) and confirmed the circularization by the detection of paired reads spanning the two ends (see below). Arrows underlining reads with the same color are paired reads (examples shown only).

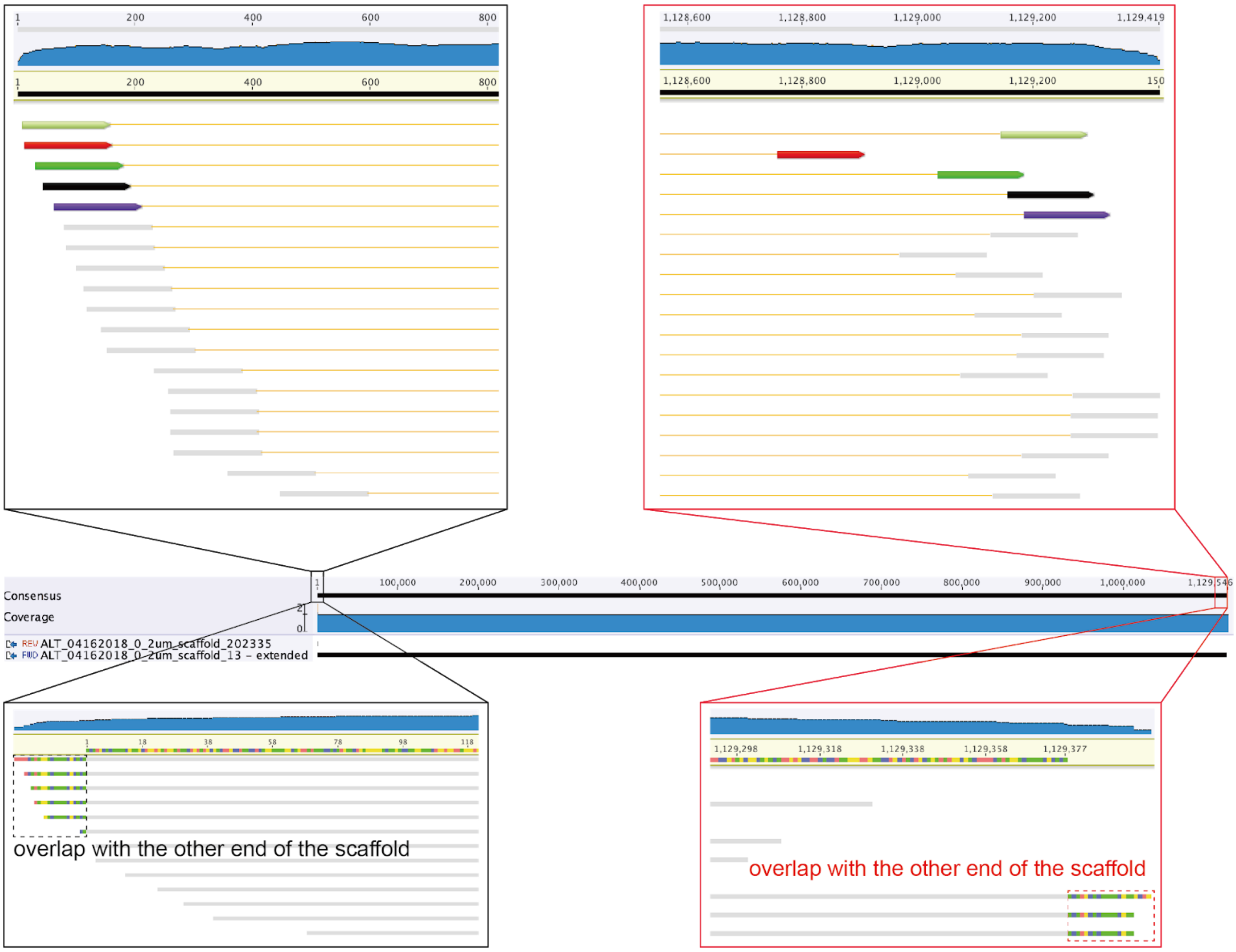

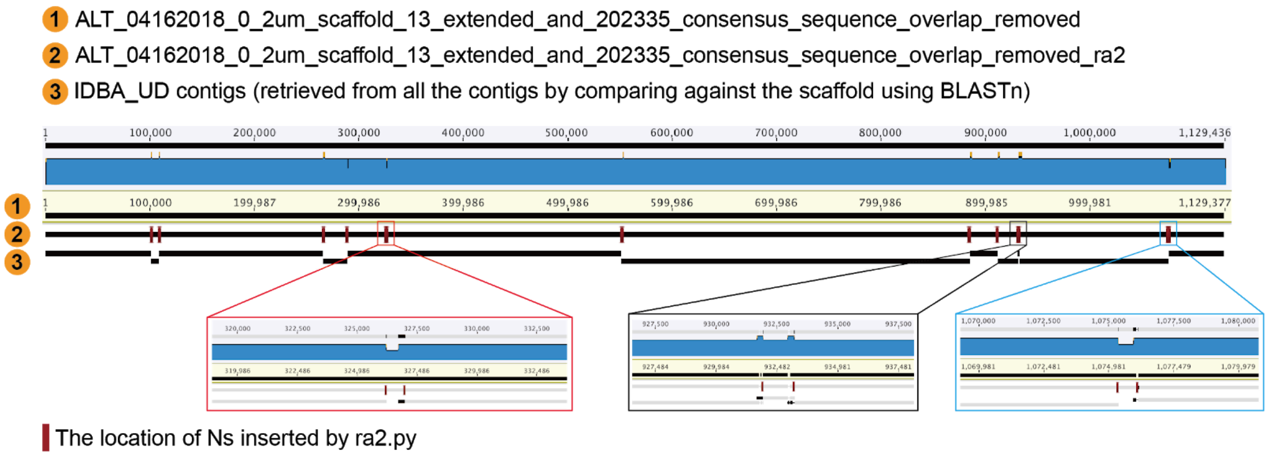


Diagram showing that local assembly errors often correspond to scaffolding joins made by the IDBA_UD assembler. 1. The scaffolds produced by IDBA_UD. 2. A total of 13 local assembly errors were reported by ra2.py (boxes). 3. The contigs generated by IDBA_UD (no scaffolding step) mapped to the scaffold. Three examples comparing the scaffolds and contigs are shown below. Errors that could not be fixed by ra2 were reported and curated manually. 12 of 13 local assembly errors were fixed as illustrated in Figures S3 and S4. The thirteenth error is in a protein-coding gene that contains multiple repeats (see below). We used the coverage of the repeat region to approximate the repeat copy number, as the problem could not be solved directly.

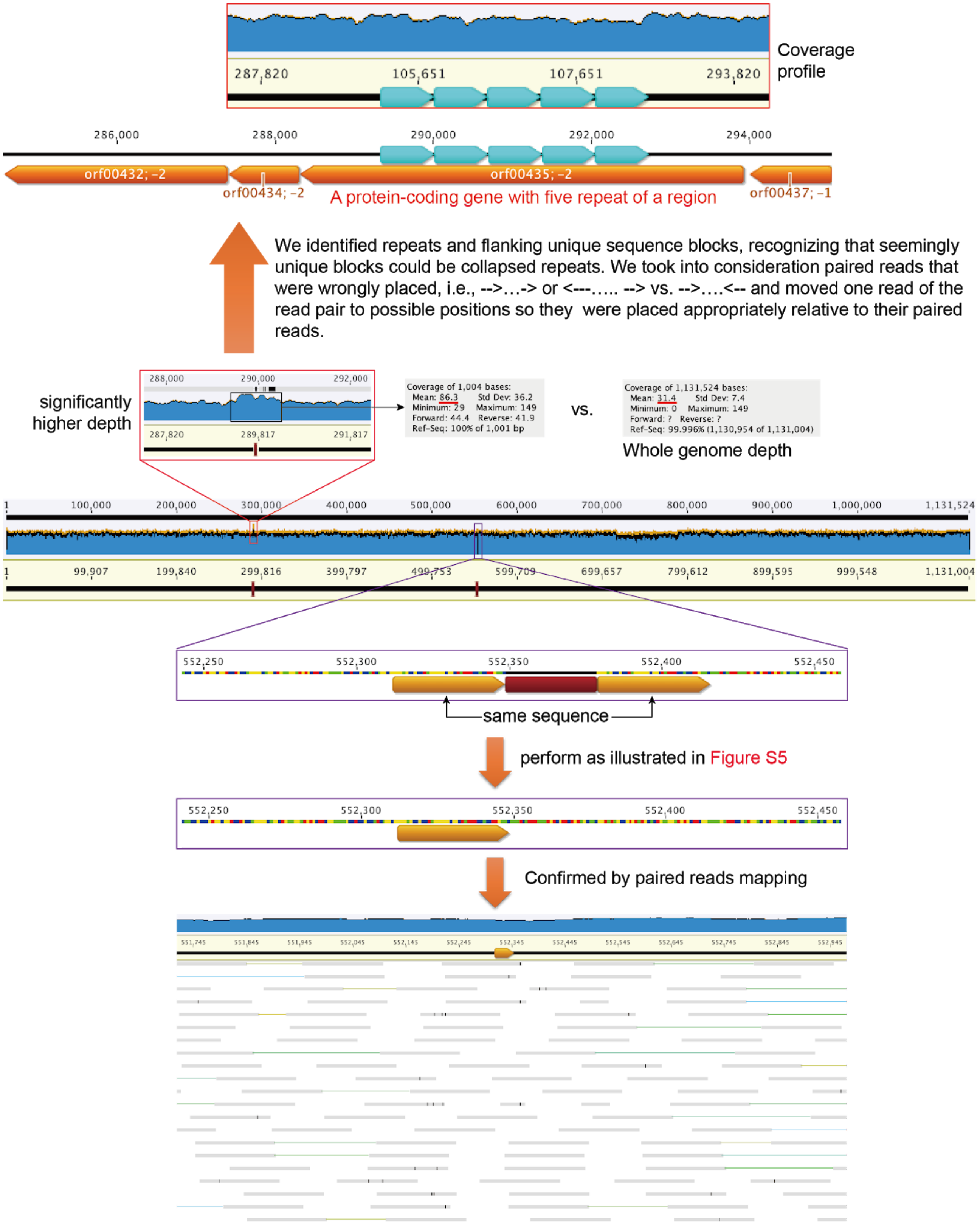
Coverage profile
287,820
105,651
107,651
293,820
286,000
288,000
290,000
292,000
294,000
orf00432; -2
orf00434; -2
orf00435; -2
orf00437; -1
A protein-coding gene with five repeat of a region
We identified repeats and flanking unique sequence blocks, recognizing that seemingly unique blocks could be collapsed repeats. We took into consideration paired reads that were wrongly placed, i.e., -->…-> or <---….. --> vs. -->….<-- and moved one read of the read pair to possible positions so they were placed appropriately relative to their paired reads.
significantly higher depth
Coverage of 1,004 bases:
Mean: 86.3 Std Dev: 36.2
Minimum: 29 Maximum: 149
Forward: 44.4 Reverse: 41.9
Ref-Seq: 100% of 1,001 bp
vs.
Coverage of 1,131,524 bases:
Mean: 31.4 Std Dev: 7.4
Minimum: 0 Maximum: 149
Forward: ? Reverse: ?
Ref-Seq: 99.996% (1,130,954 of 1,131,004)
Whole genome depth
same sequence
perform as illustrated in Figure S5
Confirmed by paired reads mapping

***Final check and adjust based on the origin of replication***: after all errors were fixed, paired reads were mapped to the genome with no mismatch allowed. As expected, all bases of the genome were well covered by reads and the coverage of the genome by reads was consistent throughout. One region had somewhat lower coverage due to the presence of prophage genome in a subset of cells. The GC skew and cumulative GC skew were calculated and the scaffold start changed to correspond with an intergenic region close to the location of the origin of replication.

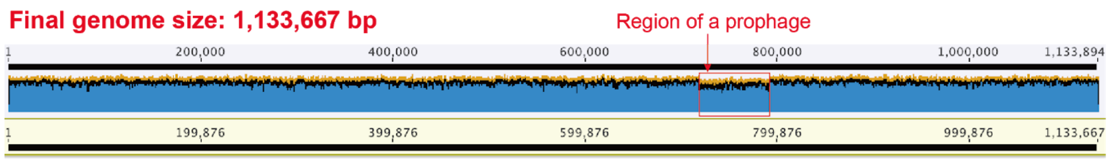


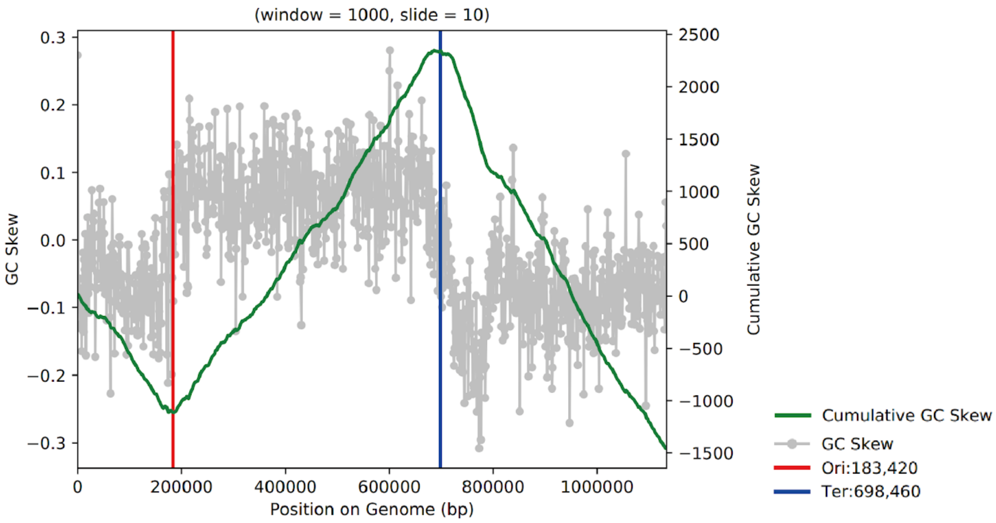


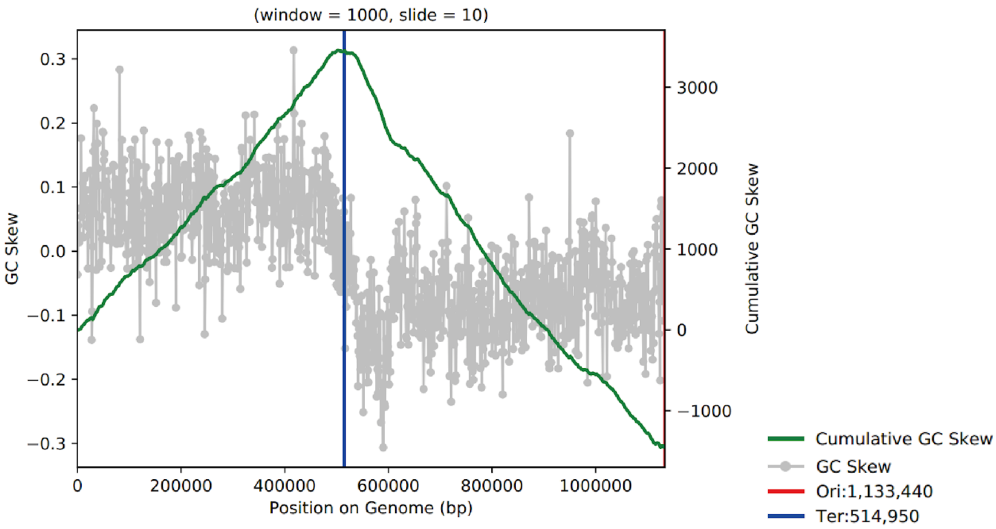

**Figure S9**. ***Case two, curation of a Betaproteobacteria genome from a MAG comprised of 7 scaffolds.***

The scaffolds were extended for the 1st run (see the extended part in blue in Figure below).

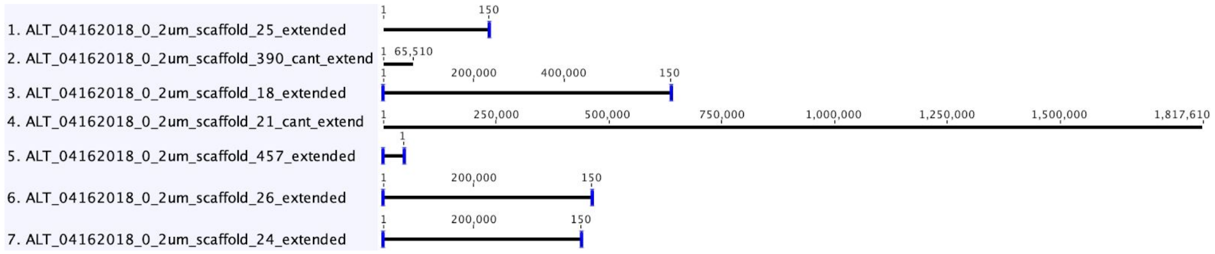


Two of the scaffolds (scaffold_21 and scaffold_25) could be assembled based on overlap and confirmed by read mapping (see below).

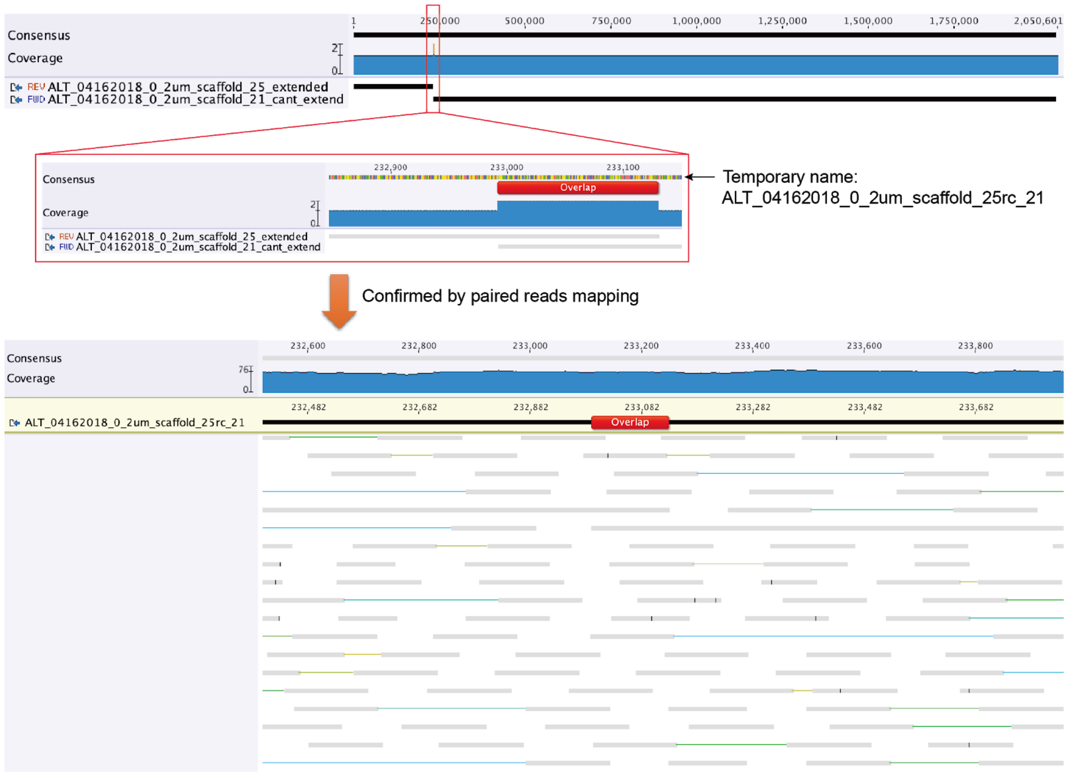


The linkage of scaffolds in the bin using fragments from the whole metagenome. Linking scaffolds were identified by BLASTn of newly extended parts of scaffolds to the full data. Possible linkages were established using "overlap-based assembly" and confirmed by reads mapping. We also considered constraints provided by the *Sulfuricella denitrificans* skB26 genome. When all the scaffolds were combined into two large genome fragments, there were two choices of how they could be arrayed. One choice, "assembled_scaffold_1 + assembled_scaffold_2" has the expected GC skew and is likely the correct solution.

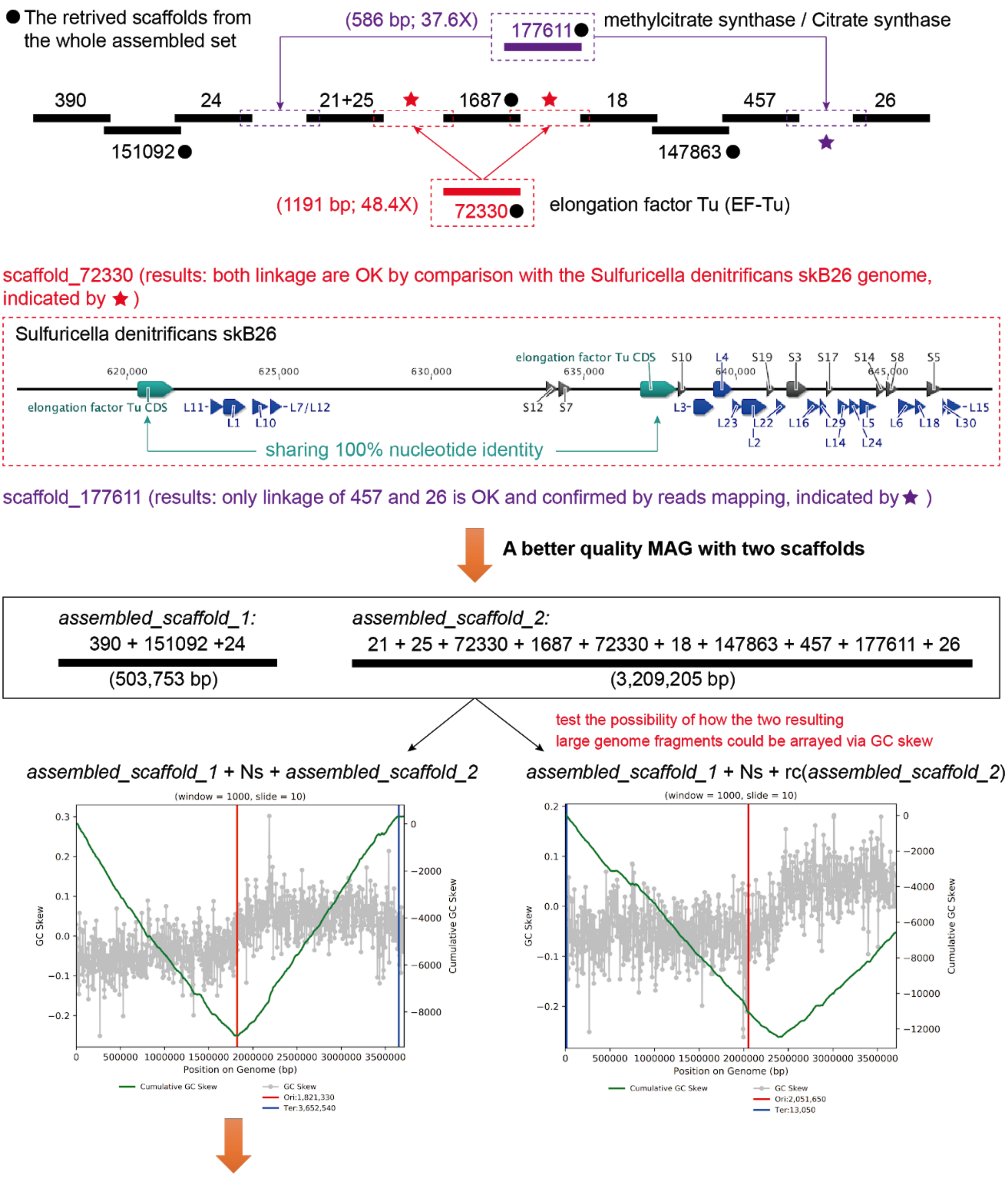


*assembled_scaffold_1* + Ns + *assembled_scaffold_2* represents the likely correct linkage of the resulting large genome fragment, the GC skew patten indicated that the genome is near complete after curation.

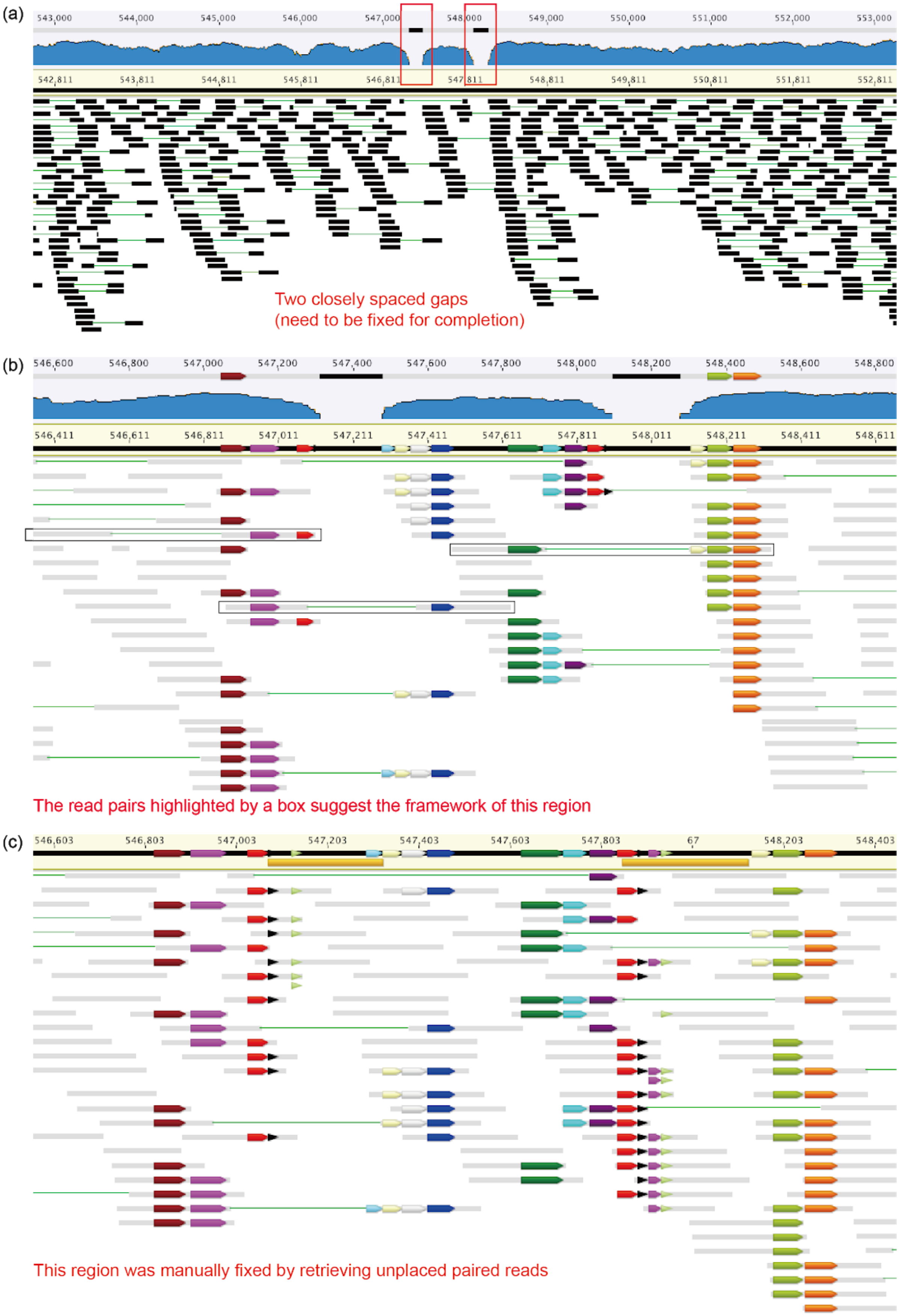


**Figure S10.** The curation of a published incomplete genome to completion was achieved by filling two closely spaced gaps. In this diagram, bars of the same length and color have the same sequence.

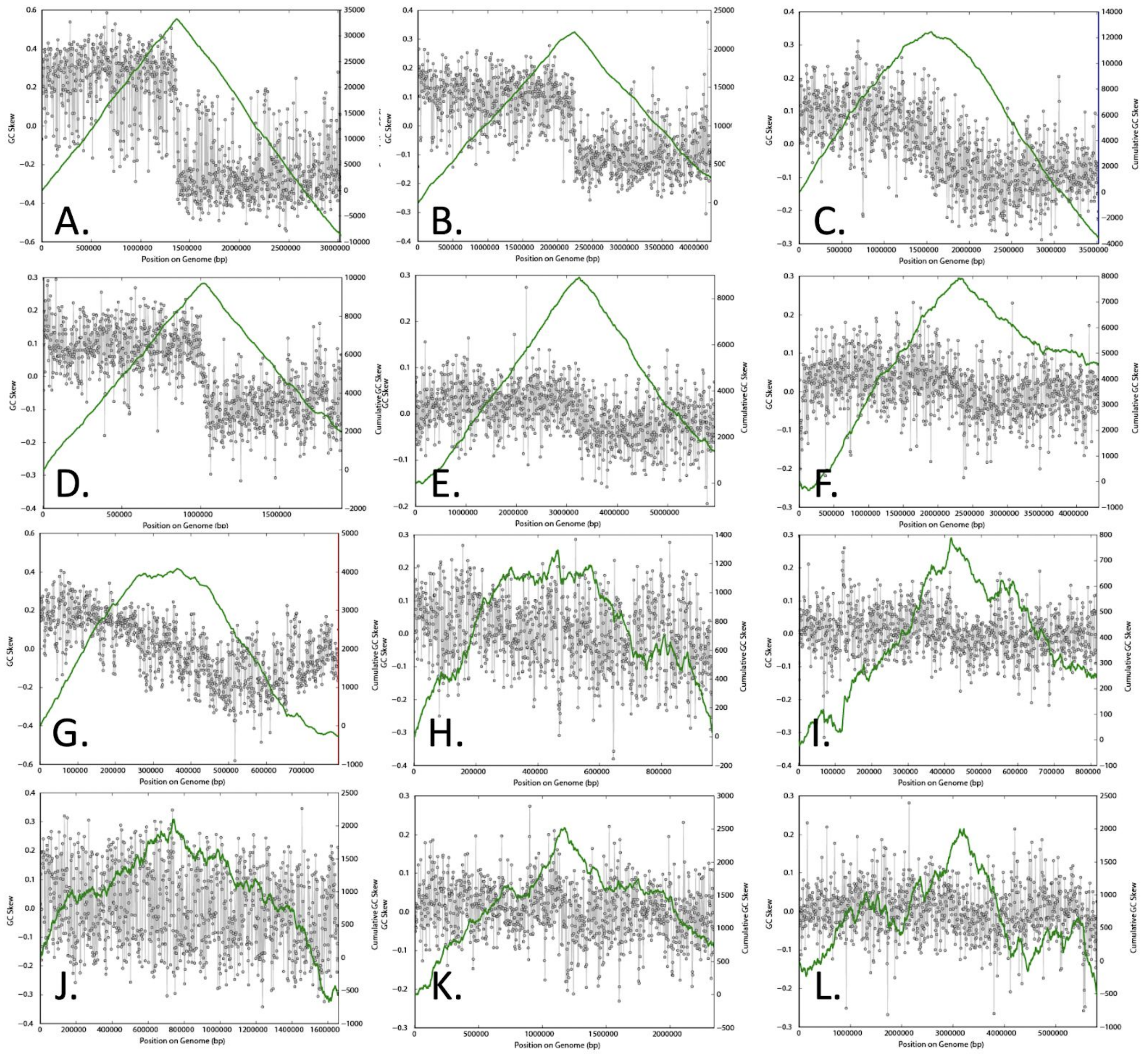


**Figure S11:** Plots of GC skew and cumulative GC skew illustrating variations in patterns across 12 complete genomes for bacterial isolates. **A.** NC_003366.1, *Clostridium perfringens* str. 13, PUBMED 11792842, **B**. NC_002570.2, *Bacillus halodurans* C-125, PUBMED 10972189, **C.** NC_006138.1, *Desulfotalea psychrophila* LSv54, DOI: 10.1111/j.1462-2920.2004.00665x, **D.** NC_004606.1, *Streptococcus pyogenes* SSI-1, PUBMED 12799345, **E.** NC_005773.3, *Pseudomonas savastanoi* pv. phaseolicola 1448A, PUBMED 16159782, **F.** NC_007802.1, *Jannaschia* sp. CCS1, no publication, **G.** NC_007292.1, *Candidatus* Blochmannia pennsylvanicus str. BPEN, PUBMED 16077009, **H.** NC_002771.1, *Mycoplasma pulmonis* UAB CTIP, PUBMED 11353084, **I.** NC_000912.1, *Mycoplasma pneumoniae* M129, DOI: 10.1093/nar/24.22.4420, **J.** NC_005072.1, *Prochlorococcus marinus* MED4, PUBMED 12917642, **K.** NC_002950.2, *Porphyromonas gingivalis* W83, PUBMED 12949112, **L.** NC_009523.1, *Roseiflexus* sp. RS-1, no publication

*Burkholderia thailandensis* E264 (NC_007651.1) (window = 1000, slide = 10)

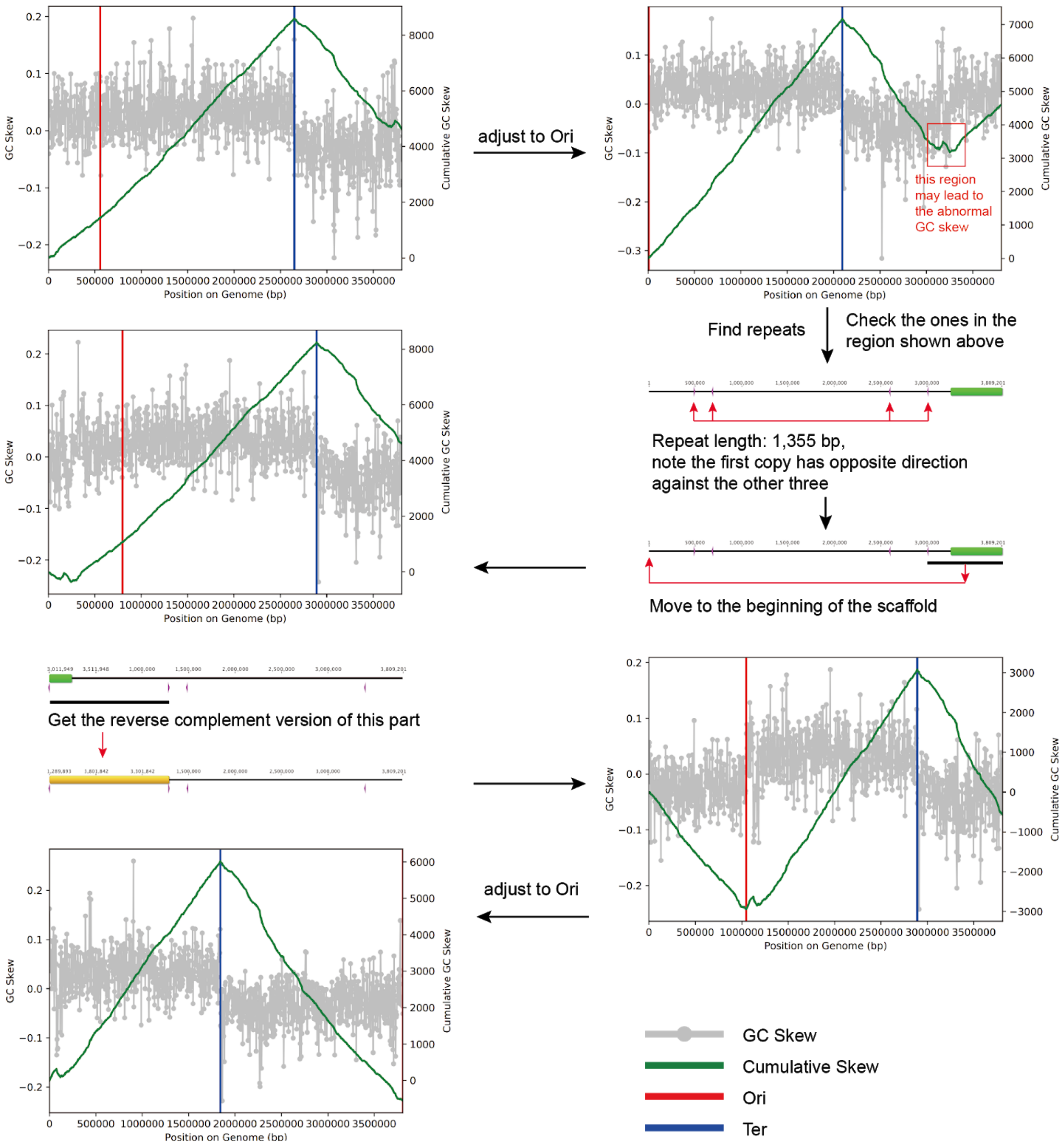


**Figure S12.** The use of GC skew and repeat analysis to identify a possible assembly error in a RefSeq genome.

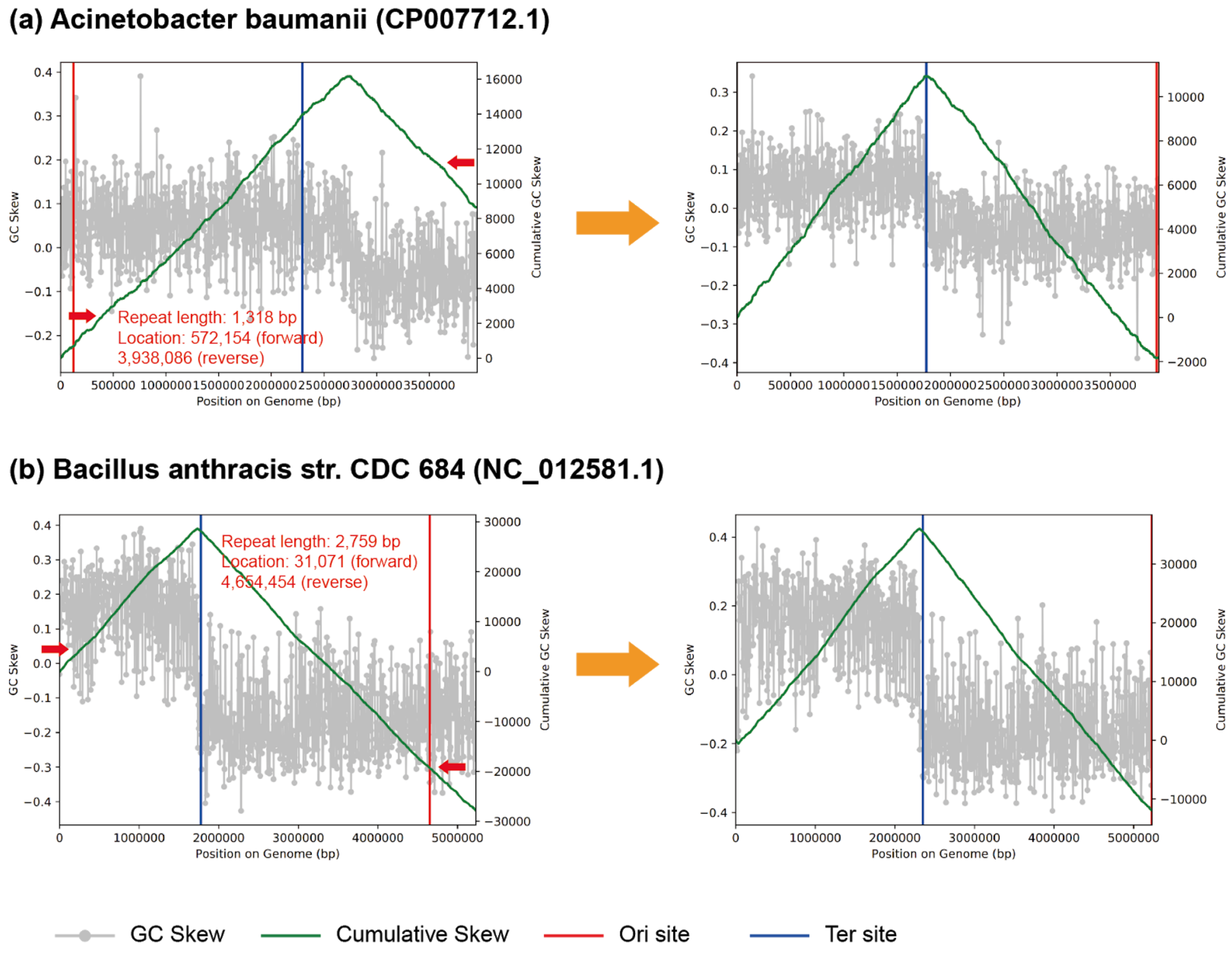


**Figure S13.** Additional examples where abnormal patterns of GC skew and cumulative GC skew suggested the presence of assembly errors in RefSeq bacterial genomes. We identified repeats (likely longer than the distance spanned by paired reads) that were in reverse complemented orientations so the intervening sequence block could be flipped and the resulting GC skew pattern profiled.
